# Novel Endogenous Engineering Platform for Robust Loading and Delivery of Functional mRNA by Extracellular Vesicles

**DOI:** 10.1101/2023.03.17.533081

**Authors:** Antje M. Zickler, Xiuming Liang, Dhanu Gupta, Doste Mamand, Mariacristina De Luca, Giulia Corso, Lorenzo Errichelli, Justin Hean, Titash Sen, Omnia M. Elsharkasy, Noriyasu Kamei, Zheyu Niu, Guannan Zhou, Houze Zhou, Samantha Roudi, Oscar P. B. Wiklander, André Görgens, Joel Z. Nordin, Virginia Castilla-Llorente, Samir EL Andaloussi

## Abstract

Messenger RNA (mRNA) has emerged as an attractive therapeutic molecule for a plethora of clinical applications. For *in vivo* functionality, mRNA therapeutics require encapsulation into effective, stable, and safe delivery systems to protect the cargo from degradation and reduce immunogenicity. Here, a bioengineering platform for efficient mRNA loading and functional delivery using bionormal nanoparticles, Extracellular Vesicles (EVs), is established by expressing a highly specific RNA-binding domain fused to CD63 in EV producer cells stably expressing the target mRNA. The additional combination with a fusogenic endosomal escape moiety, VSVg, enables functional mRNA delivery *in vivo* at doses substantially lower than currently used clinically with synthetic lipid-based nanoparticles. Importantly, the application of EVs loaded with effective cancer immunotherapy proves highly effective in an aggressive melanoma mouse model. This technology addresses substantial drawbacks currently associated with EV-based nucleic acid delivery systems and is a leap forward to clinical EV applications.

**Graphical Abstract:** (Figure created using BioRender)

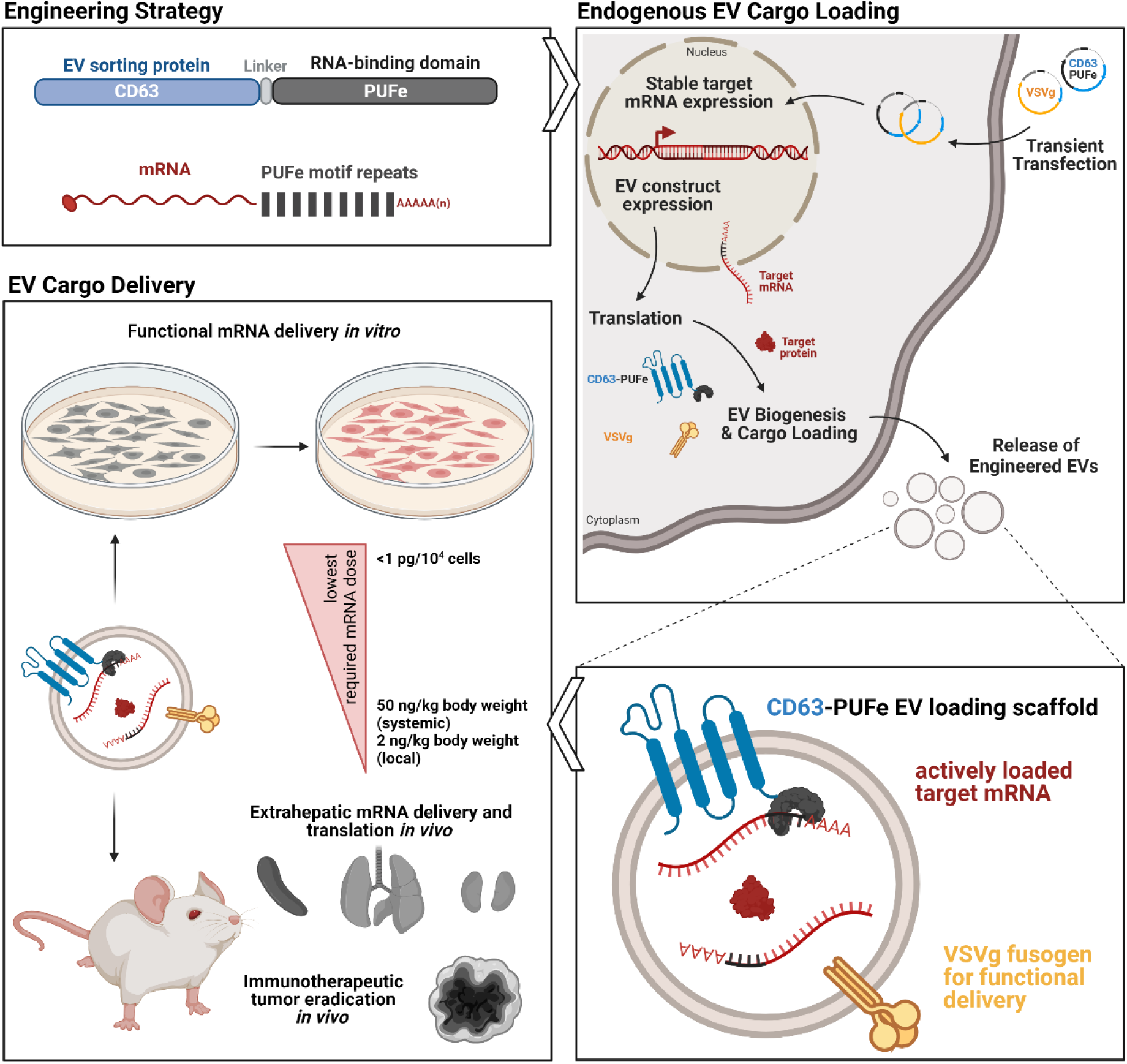

## INTRODUCTION

Nucleic acid-based therapeutics, in particular messenger RNA (mRNA), have played a pivotal role in recent developments of RNA-based vaccination platforms^1^ and cancer immunotherapy^2^. Furthermore, mRNA has emerged as an attractive molecule for the development of precision medicine approaches, such as protein-replacement therapy, gene editing, and genomic engineering^3–7^. The major limitation to the delivery of mRNA therapeutics pertains to their instability and inherent inability to bypass the lipid bilayer of cells^8^. Current state-of-the-art for therapeutic mRNA delivery is the encapsulation into synthetic nanoparticles, such as lipid nanoparticles (LNPs). Although their efficacy is indisputable for the delivery of mRNA-based SARS-CoV-2 vaccines for the prevention of coronavirus disease 2019 (COVID-19)^9–11^, their clinical applications are sometimes associated with adverse effects of different severities, including immunogenicity due to their synthetic nature^12–14^. Moreover, targeted RNA delivery to distant organs is challenging using LNPs^8,15^.

An alternative nanotechnology-based approach is the use of engineered extracellular vesicles (EVs), including exosomes, for the delivery of mRNA therapeutics. EVs are lipid bilayer-enclosed, naturally secreted nanovesicles of 30-2000 nm in size^16^, are immunologically tolerated and can cross biological barriers to distant organs^17,18^. They contain macromolecular cargo from the source cell and can deliver these proteins, nucleic acids, or lipids to recipient cells^17,18^. Most importantly, they can be bioengineered to carry luminal therapeutic cargo as well as target molecules on their surface^17,19–23^.

Incorporation of comparatively large mRNA cargo into pre-isolated EVs using exogenous loading approaches, such as co-incubation or electroporation, is cumbersome and quite inefficient, and thus not sustainable in a clinical setting^24–26^. In contrast, endogenous loading approaches involving genetic engineering of the parental cells to incorporate mRNA cargo during EV biogenesis, are much more promising for therapeutic application. However, specific enrichment of mRNA cargo in EVs proves inefficient without an active loading strategy^27^. Thus, cells are typically engineered with genetic constructs expressing an EV sorting protein^22^ fused to an RNA binding domain (RBD), and the mRNA of interest. Correspondingly, RBD-interacting sites are inserted in the UTRs of the cargo mRNA (Targeted and Modular EV Loading (TAMEL) approach)^28–33^.

Substantial drawbacks of current endogenous EV engineering strategies for mRNA loading include insufficient mRNA loading despite active EV sorting and insufficient cargo delivery to target cells. Moreover, unsolicited carry-over of plasmid DNA potentially interferes with the assessment of mRNA quantification and functionality^34–36^. To address these shortcomings, we developed a novel TAMEL-based platform for efficient mRNA delivery using engineered EVs. The target mRNA expression cassette was stably integrated into the genome of EV producer cells to increase assay robustness and avoid mRNA-encoding plasmid carry-over. As EV sorting scaffold CD63, a tetraspanin abundantly present on EVs, was fused to the high-affinity mammalian RBD PUFe, an optimized version of the designer Pumilio and FBF homology domain^37^. The engineered EVs demonstrated efficient, robust, and reproducible mRNA cargo loading, and functional mRNA delivery *in vitro*. To enhance delivery efficiencies by induced endosomal escape, the EV producer cells were further engineered to express the viral fusogen Vesicular Stomatitis Virus Glycoprotein (VSVg). With this strategy the engineered EVs showed extrahepatic mRNA delivery upon systemic injection *in vivo* at doses as low as 50 ng mRNA per kg body weight. Importantly, our EV engineering platform robustly achieved complete remission in up to 67% of model mice with highly aggressive melanoma when engineered EVs were loaded with the immunostimulatory co-receptor Ox40L mRNA and injected intratumorally.

## RESULTS

### Generation of an EV-specific mRNA loading platform

To realize specific endogenous loading of engineered mRNA into EVs, a two-component system was designed. The approach combined an EV loading scaffold, i.e. an EV sorting domain fused to an RBD (Figure 1a), with a co-expressed target mRNA containing multiple repeats of the RBD-dependent high-affinity motif in the 3’UTR (Figure 1b). Here, the tetraspanin CD63, was fused to either the non-cleaving S148A mutant of the CRISPR-associated protein Cas6f^38^, or three engineered RBDs derived from the designer Pumilio and FBF homology domain (PUF)^37^, PUFm, PUFe and PUFx2, respectively (Figure 1a). PUFm was chosen for its high affinity to the RNA target sequence (K_d_ ≈ 4 pM)^39^, PUFe was engineered to bind 8 nucleotides of low abundance in mammalian cells rendering it highly specific for the target RNA^37^, and PUFx2 was designed to bind to a longer stretch of 16 target nucleotides^40^. Correspondingly, mRNA expression constructs were engineered with 6x 28-nt hairpin structures for Cas6f (S148A) binding and 10x high-affinity motifs for binding to the respective PUF-derived RBD (Figure 1b). The mRNA expression constructs were designed for Nanoluciferase (Nanoluc)^41^ and Cre recombinase (Cre) reporter proteins, as well as murine Ox40L as a therapeutic protein. To account for the passive loading of mRNA or protein into EVs, CD63 was also fused to the bacteriophage MS2 coat protein as an incompatible RBD not binding to the target mRNA, termed here “non-binding control” (NBC, Figure 1a).

**Figure 1.**
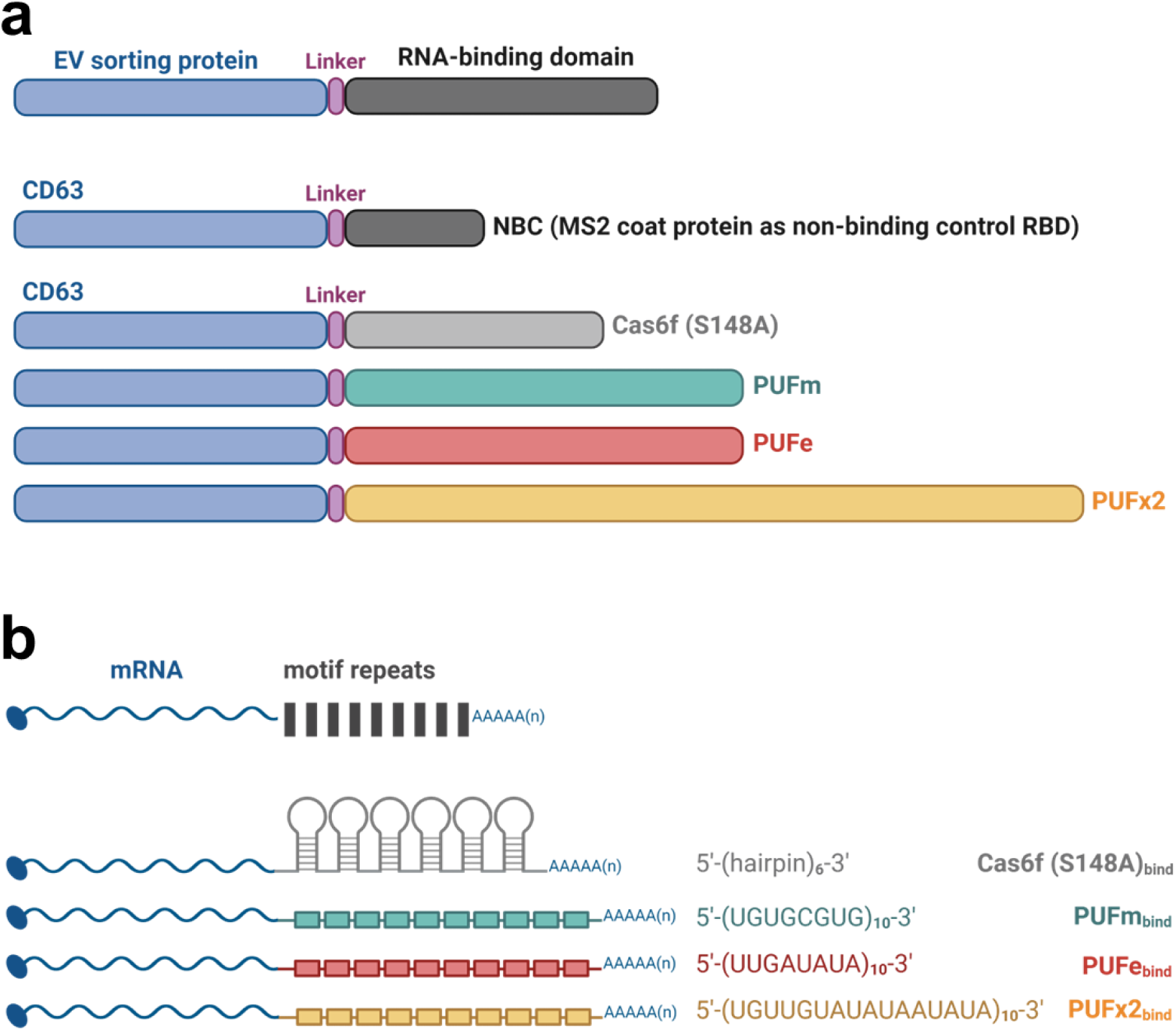
Engineering strategy for endogenous mRNA loading into EVs. (a) EV sorting strategy: four novel RNA-binding domains (RBD), the non-cleaving mutant Cas6f and three engineered versions of the designer Pumilio and FBF homology domain (PUF), termed PUFm, PUFe, and PUFx2, were fused via a glycine-rich linker peptide to the C-terminus of the EV sorting protein CD63. As control for passive loading (non-binding control, NBC), CD63 was fused to the MS2 coat protein (MCP, as published by Hung et al., 2016). (b) mRNA-binding strategy: mRNA coding sequences were codon-optimized and engineered to contain 6-10 repeats of the specific high-affinity motif recognized by the respective RNA-binding domain in their 3’UTR. Figure was created using BioRender.

### The CD63-PUFe EV scaffold efficiently loads Nanoluc mRNA into the lumen of engineered EVs

Endogenously engineered EVs for therapeutic mRNA loading are often produced by transient co-transfection of multiple plasmids to overexpress engineering constructs in EV producer cells^28,42,43^ . This approach has proven effective, yet awareness of the potential pitfalls, such as residual plasmid or transfection reagent in the EV preparation, is required when interpreting experimental results from EV cargo quantification and delivery assays^34–36^. Additionally, the inclusion of appropriate controls to account for the passive loading of both target mRNA and the resulting protein is crucial for interpreting the loading and delivery correctly^27^. Here, HEK293T EV producer cells were engineered to stably express the target mRNA from a genomic locus using the ϕC31 integrase system^44,45^ (Figure 2a). Stable mRNA-expressing producer cells were subsequently transfected with plasmids encoding for the EV loading construct for EV production. This approach allowed for the generation of EV preparations without mRNA-encoding plasmid and ensured similar rates of passive loading. To evaluate mRNA loading efficiencies, EVs that were both actively loaded (target mRNA EVs) and passively loaded (target control EVs) were generated from the same stable cell line. CD63-NBC (non-binding control) was used as control for passive loading in all experiments. With this approach we ensured that both mRNA EVs and control EVs derived from the same engineered producer cells.

**Figure 2.**
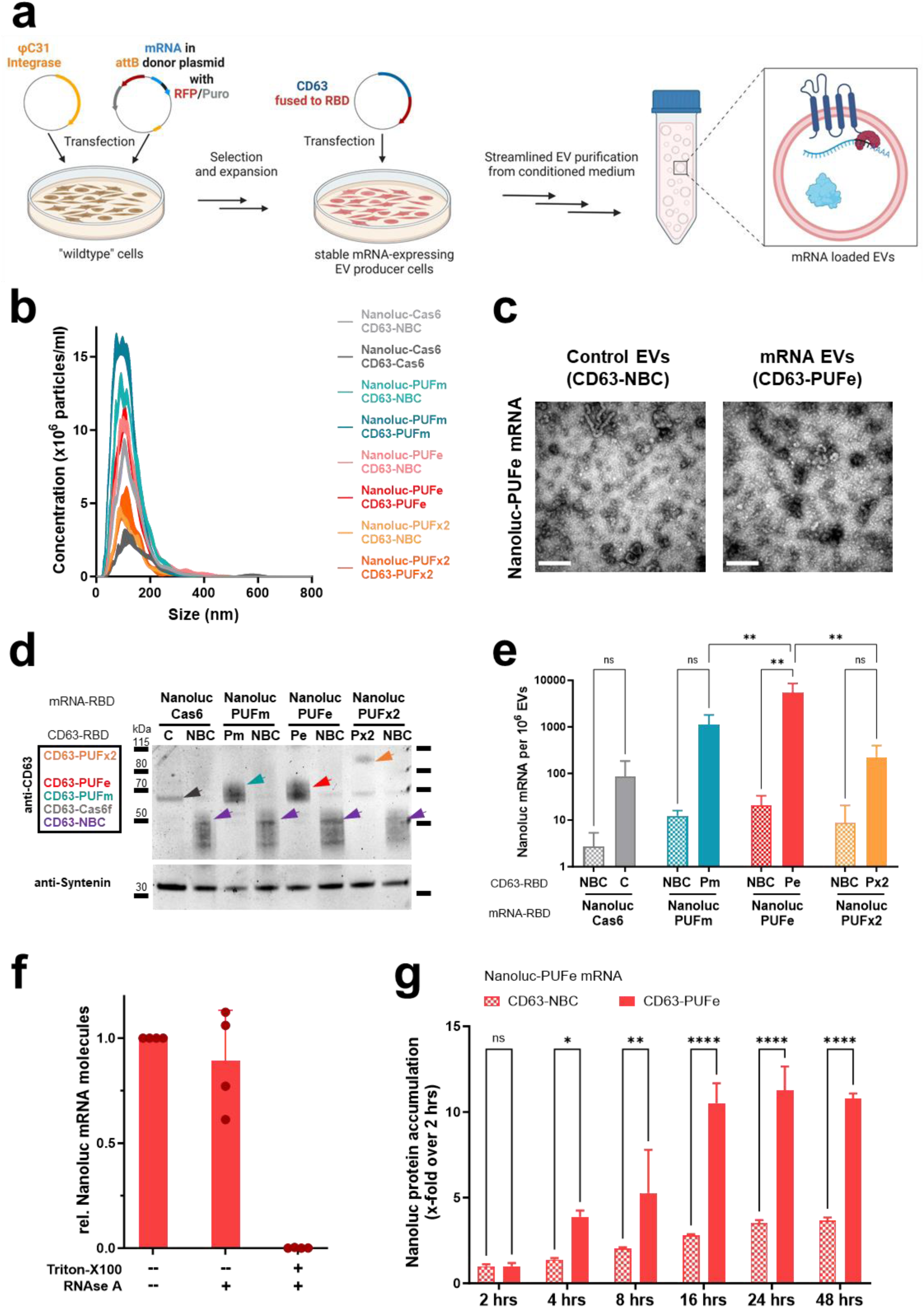
Active mRNA loading into EVs from engineered mRNA single-stable producer cells. (a) Illustration showing mRNA stable cell line generation using the ϕC31 integrase system, and subsequent EV production. For both mRNA EV and control EV production, the same mRNA stable EV producer cell line was used. Cells were either transiently expressing the compatible CD63-RBD or incompatible CD63-NBC (non-binding control). This approach ensured the same biological pre-requisites during EV biogenesis for passive loading of engineered mRNA and protein. Figure created using BioRender. (b) Average particle size determination by NTA of mRNA EVs and respective control EVs harvested from transfected Nanoluc-RBD mRNA stable producer cells (RBD motif as indicated). (c) Negative stain Transmission Electron Microscopy images of control EVs and mRNA EVs loaded with Nanoluc-PUFe mRNA. Scale bar: 300 nm (d) Western Blot analysis of EVs produced from Nanoluc-RBD mRNA stable EV producer cells as indicated on top. The expression of the CD63-RBD fusion proteins, either CD63-Cas6f (C), CD63-PUFm (Pm), CD63-PUFe (Pe), CD63-PUFx2 (Px2), or CD63-NBC (NBC), respectively, was validated by probing for CD63, sizes are indicated with colored arrowheads. As loading reference, expression of the EV marker SDCBP (Syntenin) was detected. Protein loaded per lane: 3 µg (e) Absolute quantification by RT-qPCR of Nanoluc mRNA molecules per 1×10^6^ control EVs (NBC) or mRNA EVs (C, Pm, Pe, Px2) averaged from three representative experiments. CD63-PUFe repeatedly showed significant enrichment of Nanoluc mRNA in EVs. (f) RNAse challenge assay to determine the efficiency of intraluminal EV mRNA cargo encapsulation in 4 independently produced EV batches. (g) Uptake of Nanoluc mRNA EVs at a dose of 0.6 pg Nanoluc mRNA per 1×10^4^ cells or particle count-matched control EVs in Huh7 recipient cells. Cellular Nanoluc protein activity was measured at indicated timepoints and normalized to the signal measured at 2 hrs, which corresponds to the signal from passively loaded Nanoluc protein. Uptake of Nanoluc mRNA EVs led to a significantly higher Nanoluc protein accumulation over time compared to control EVs, demonstrating EV-mediated engineered mRNA delivery and functional translation.

After EV isolation, the average particle diameter was verified by NTA at 101 (± 2.6) nm (Figure 2b) and the presence of EVs was confirmed by electron microscopy (Figure 2c, scale bar 300 nm). Next, CD63-RBD constructs and reference protein expression in both EVs and producer cells were validated by Western Blotting (Figure 2d and Supplementary Figure S 1a, respectively). Nanoluc protein abundance was determined by luminescence measurements, which showed that the engineered HEK293T cell lines contained comparable amounts of Nanoluc protein stemming from equivalent copy numbers of genomically integrated mRNA expression cassettes (Supplementary Figure S 1b). The purified mRNA EVs and control EVs also carried similar quantities of passively loaded Nanoluc protein (Supplementary Figure S 1c).

Next, the mRNA loading efficiencies for the four novel CD63-RBD loading scaffolds were assessed. While the relative expression of Nanoluc mRNA in producer cells was similar in all samples with a maximum difference of 1.5-fold (Supplementary Figure S 1d), the actively loaded mRNA EVs contained substantially more Nanoluc mRNA molecules as determined by absolute quantification using RT-qPCR (Figure 2e). The enrichment over control values ranged from 18.5-fold for CD63-Cas6, to 65-fold for both PUFm and PUFx2, and reached more than 230-fold for CD63-PUFe (Figure 2e). To evaluate EV integrity and cargo encapsulation efficiency, intraluminal loading of mRNA for CD63-PUFe mRNA-loaded EVs was validated by an RNAse challenge assay. The majority of mRNA, 89% on average in four tested EV batches, was loaded within the EV lipid bilayer and was therefore protected from RNAse degradation (Figure 2f).

In summary, we established a novel engineering approach for EV mRNA loading using stable target mRNA expression in EV producer cells, where CD63-PUFe proved to function as the most robust and efficient mRNA loading scaffold for EV bioengineering and was thus selected for all subsequent experiments to establish an EV-mediated mRNA delivery system.

### CD63-PUFe engineered EVs functionally deliver Nanoluc mRNA *in vitro*

To assess the translatability of EV-derived Nanoluc-PUFe mRNA purified EV RNA was transfected into Huh7 and HEK293T recipient cells and Nanoluc protein activity was measured at different time points (Supplementary Figure S 2a and S 2b). While RNA from control EVs only yielded minor Nanoluc protein expression in transfected cells, RNA from Nanoluc-PUFe mRNA EVs showed substantially enhanced translation of Nanoluc protein in both cell lines at all timepoints, proving functionality of EV-derived mRNA. Next, EV uptake experiments *in vitro* were performed to evaluate the release and translation of EV-encapsulated mRNA in recipient cells. First, Huh7 cells were treated with escalating doses of EVs actively loaded with Nanoluc-PUFe mRNA or the particle-matched amount of passively loaded control EVs (Supplementary Figure S 2c). After 24 h, treated cells were analyzed for Nanoluc protein activity. To account for Nanoluc protein not derived from mRNA translation, the luminescence from EV-treated cell lysate in this and all subsequent Nanoluc mRNA EV uptake experiments was normalized to the respective EV luminescence, the total input of passively loaded Nanoluc protein before translation (exemplified in Supplementary Figure S 2d). For all three tested EV-encapsulated Nanoluc-PUFe mRNA doses (0.02 – 2 pg per 10^4^ cells), we observed minor, but insignificant, increases in ratios of Nanoluc protein expression in cells treated with Nanoluc-PUFe mRNA EVs (Supplementary Figure S 2c). To identify the optimal time frame for EV-mediated mRNA delivery in EV uptake assays, the uptake of Nanoluc-PUFe mRNA EVs and control EVs was tested over time. Here, the accumulation of Nanoluc protein in the recipient Huh7 cells was measured relative to the 2-hours’ time point under the assumption that Nanoluc-PUFe mRNA translation has not started, while delivered Nanoluc protein can already be detected. Nanoluc protein accumulation in cells treated with mRNA EVs reached its peak at 24 hours (11.2-fold increase over 2 hours) of EV incubation (Figure 2g). In contrast, cells treated with particle-matched control EVs showed a slow accumulation of EV protein plateauing at 24 hours (3.5-fold increase over 2 hours). In comparison to control, the overall Nanoluc protein accumulation was significant in cells treated with mRNA EVs. This data indicated an active supply of newly translated Nanoluc protein from delivered mRNA in cells treated with Nanoluc-PUFe mRNA EVs.

### VSVg-mediated endosomal escape is crucial for enhanced EV cargo delivery *in vitro* **and *in vivo***

Despite the encouraging EV uptake results *in vitro*, the rate of Nanoluc protein expression upon EV-mediated mRNA delivery did not reflect the significant enrichment of mRNA in actively loaded EVs. This observation is likely due to inefficient endosomal escape and thus premature cargo degradation in the target cells after EV uptake. The EV producer cells were therefore engineered to co-express the fusogenic protein VSVg, previously shown to facilitate the delivery of EV cargo by inducing endosomal escape in the target cell^33,46,47^.

To assess the impact of VSVg expression on loading and EV-mediated mRNA delivery, Nanoluc-PUFe mRNA EVs and control EVs were produced without (mock) or with VSVg co-expression (Figure 3a). The presence of EVs was validated by NTA with a mean peak diameter at 112.9 ± 9.5 nm (Figure 3b), and the expression of CD63, VSVg, and Nanoluc protein were validated by Western Blot in both EVs and producer cells (Figure 3c). The EV mRNA loading efficiencies were determined by RT-qPCR and showed more efficient mRNA loading in absolute numbers for Nanoluc-PUFe mRNA EVs without VSVg (average 2217 molecules per 10^6^ EVs) as compared to Nanoluc-PUFe mRNA EVs carrying VSVg (average 1459 molecules per 10^6^ EVs), while the enrichment over control was similar for both, 410-fold and 430-fold, respectively (Figure 3d).

**Figure 3.**
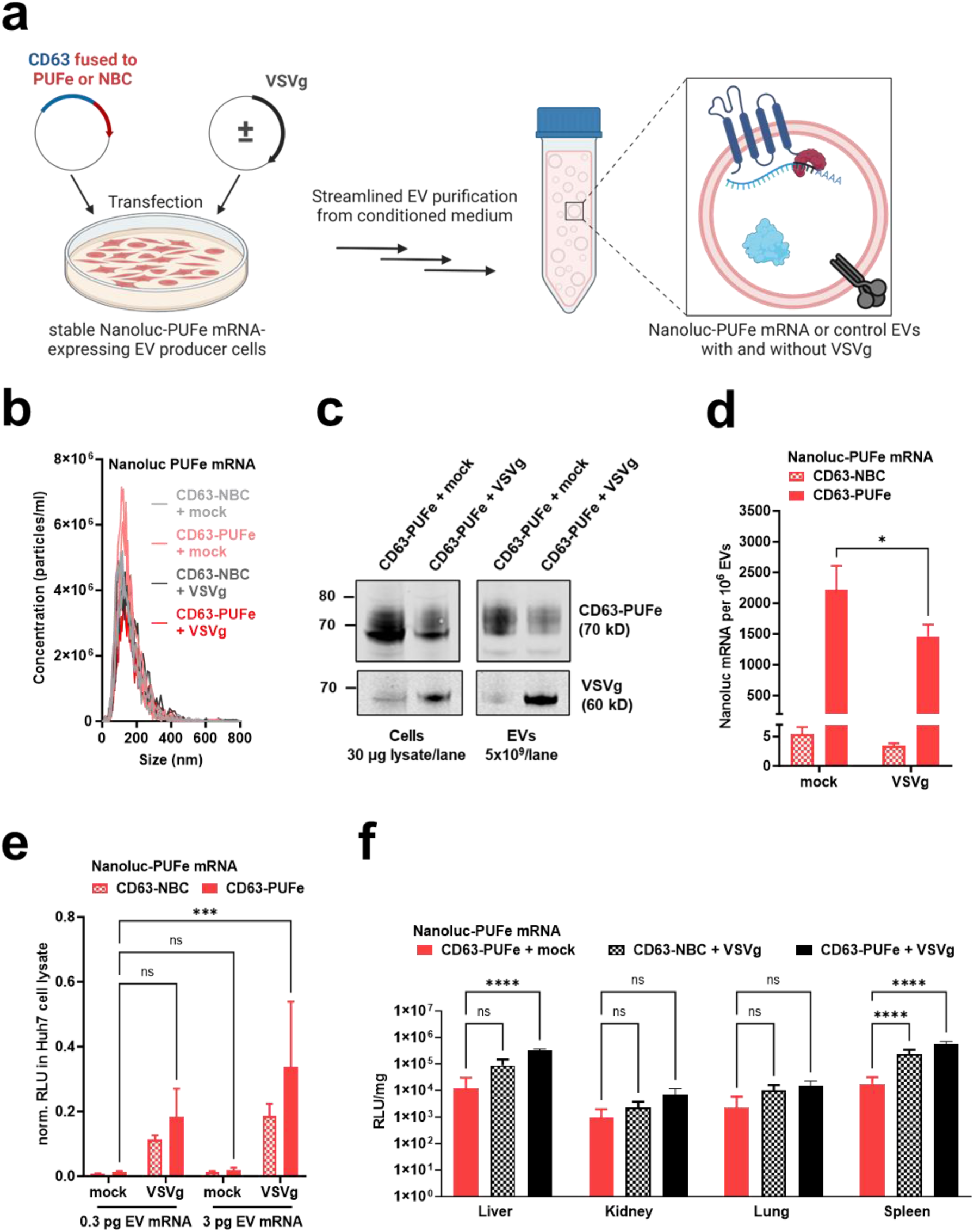
Co-expression of the viral fusogen VSVg induces endosomal escape and substantially increased EV cargo delivery. (a) Illustration of VSVg co-expression in Nanoluc-PUFe mRNA stable EV producer cells, and subsequent EV production. Figure created using BioRender. (b) Average particle size determination by NTA of Nanoluc-PUFe mRNA EVs and control EVs with or without VSVg co-expression. (c) Western Blot analysis to validate CD63-PUFe and VSVg expression in Nanoluc-PUFe mRNA EV producer cells (left) and corresponding EVs (right). (d) Absolute quantification by RT-qPCR of Nanoluc-PUFe mRNA loading into EVs with or without VSVg co-expression. (e) *In vitro* uptake analysis of Huh7 recipient cells treated with equal amounts of Nanoluc-PUFe mRNA EVs at a dose of0.3 pg and 3 pg mRNA per 1×10^4^ cells, or particle count-matched control EVs, both with and without VSVg expression. Co-expression of VSVg on the EVs significantly enhanced Nanoluc protein signal in treated cells after 24 h, indicating a general improvement of EV cargo release by VSVg-mediated endosomal escape. (f) Co-expression of VSVg enhanced EV cargo delivery *in vivo*. Mice were injected intraperitoneally with Nanoluc mRNA EVs with or without VSVg co-expression at a Nanoluc mRNA dose of 50 ng/kg body weight, or particle count-matched control EVs with VSVg co-expression (n=4 per group). Nanoluc protein signal was measured in the liver and extrahepatic organs at 24 hours post injection and showed extrahepatic mRNA delivery as well as higher protein expression in mice treated with EVs co-expressing VSVg.

Next, the impact of VSVg in EV-mediated mRNA delivery was assessed in uptake assays *in vitro* over 24 h where Huh7 cells were treated with two doses of EVs according to their Nanoluc-PUFe mRNA content. EVs displaying VSVg showed significantly higher efficiencies in dose-dependent cargo delivery of both Nanoluc protein and mRNA, as compared to mRNA EVs without VSVg (Figure 3e).

To substantiate our findings *in vivo*, Nanoluc-PUFe mRNA EVs without VSVg and with VSVg were injected intraperitoneally (ip) into mice at a Nanoluc mRNA dose of 50 ng/kg bodyweight, in comparison to particle count-matched control EVs with VSVg. At 24 h after injection organs were harvested and analyzed for Nanoluc protein expression (Figure 3f). A generally elevated Nanoluc protein abundance was observed in all analyzed organs when VSVg was present, irrespective of the amount of loaded mRNA. However, Nanoluc activity in mice treated with mRNA EVs carrying VSVg exceeded the values of control EV-treated mice, showing successful EV-mediated mRNA delivery and translation in liver and extrahepatic organs in the presence of VSVg (Figure 3f).

Taken together, our results clearly demonstrated the key role of endosomal escape in EV-mediated cargo delivery, as the display of VSVg as fusogenic protein proved crucial for Nanoluc mRNA and protein delivery *in vitro*, while also significantly enhancing extrahepatic delivery *in vivo*.

### The CD63-PUFe EV mRNA loading platform efficiently delivers gene editing modalities

Although not problematic from a therapeutic perspective, in our described system the highly sensitive detection of co-delivered passively loaded Nanoluc protein may strongly mask the true signal from newly translated protein derived from the delivered Nanoluc-PUFe mRNA. To ultimately prove the functionality of EV-delivered mRNA we tested the platform by delivering Cre recombinase mRNA to Cre reporter cells *in vitro*. We attempted the generation of Cre- PUFe mRNA stable cells, however, the integrated genomic cassette was repeatedly silenced and Cre mRNA expression was declining rapidly during the first passages after selection (data not shown). Therefore, Cre-PUFe mRNA and control EVs were produced by triple co-transfection from wildtype HEK293T cells (Figure 4a).

**Figure 4.**
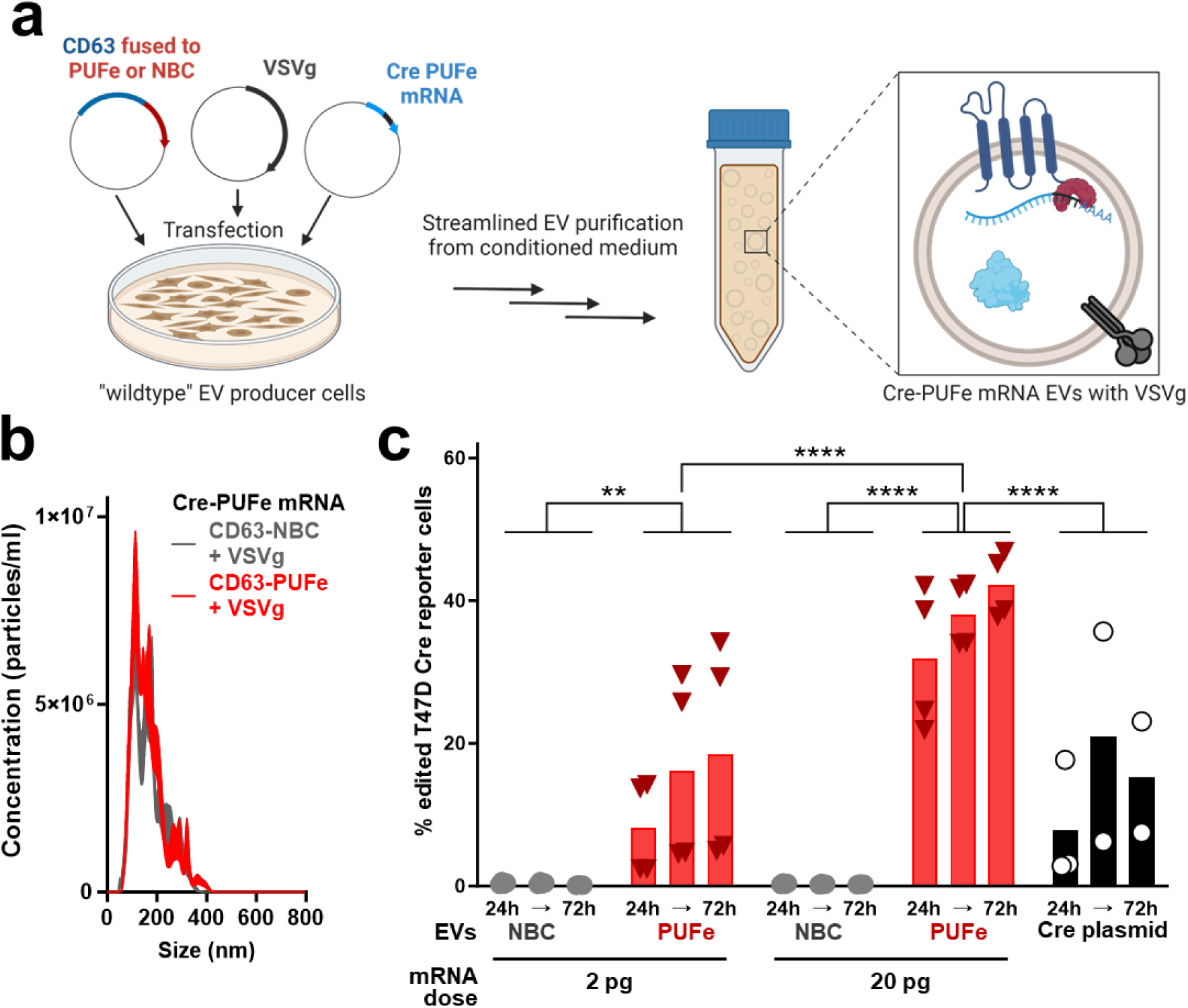
Engineered EVs efficiently delivered Cre recombinase mRNA *in vitro*. (a) Illustration of triple co-transfection methodology for Cre-PUFe mRNA EV production from HEK293T producer cells, and subsequent EV production. Figure created using BioRender. (b) Average particle size determination by NTA of Cre-PUFe mRNA EVs and control EVs with VSVg co-expression. (c) *In vitro* uptake analysis of T47D Traffic Light Cre reporter cells treated with equal amounts of Cre-PUFe mRNA EVs at a dose of 2 pg and 20 pg mRNA per 1×10^4^ cells, or particle count-matched control EVs, both with VSVg expression. Frequencies of genomically edited Cre reporter cells were assessed at 24, 48, and 72 hours post treatment by flow cytometry. Cells treated with mRNA loaded EVs show a time-and dose-dependent increase in genomically edited cell frequencies, even exceeding efficiencies achieved by Cre plasmid transfection (black). Significance level α=0.05, Two-way ANOVA.

After EV production and particle size validation (137.4 ± 19.5 nm at peak concentration, Figure 4b), loaded Cre mRNA molecules in EVs were measured. Again, the active loading platform enriched mRNA EVs substantially for Cre-PUFe mRNA compared to control EVs (Supplementary Figure S 3a).

To test mRNA delivery functionality, B16F10 and T47D Traffic Light Cre reporter cells were treated with escalating doses of Cre-PUFe mRNA or particle count-matched control EVs. Here, significant frequencies of genomically recombined T47D reporter cells was detected in cells treated with mRNA EVs over all time points and mRNA doses (Figure 4c). For B16F10 Cre reporter cells, significant frequency of recombination was observed only for cells treated with the higher dose of EVs (Supplementary Figure S 3b). Additionally, at the highest dose of mRNA EVs, the recombination efficiencies in T47D Cre reporter cells exerted by EV-delivered mRNA outperformed those observed in Cre plasmid-transfected cells, highlighting the high efficiency of our system (Figure 4c). The negligible recombination in the control group furthermore ruled out delivery of co-purified plasmid DNA, passively loaded Cre-PUFe mRNA, or passively loaded Cre protein, corroborating earlier studies on Cre protein loading into EVs^47^.

Next, we benchmarked the performance of the CD63-PUFe-based EV mRNA delivery platform against the selective endogenous encapsidation for cellular delivery (SEND)^30^ technology. EVs actively loaded with Cre mRNA or control EVs were produced using our EV loading scaffold or the human PEG10-based SEND platform. The expression of key proteins in EVs and producer cells were verified by Western Blot (Supplementary Figure S 3c). B16F10 Traffic Light Cre reporter cells were incubated with 20 µl purified mRNA-loaded or control EV suspension, and genomic recombination was assessed at 24 h, 48 h, and 72 h, as published^30^. Cre mRNA delivery using the CD63-PUFe platform was highly efficient at all three analyzed time points, while recombination efficiencies were lower for mRNA-loaded EVs produced with the SEND packaging technology (Supplementary Figure S 3d).

In summary, the CD63-PUFe EV mRNA loading platform performed as an advanced application of the TAMEL technology for high-efficiency EV-mediated delivery of gene editing mRNA modalities *in vitro*.

### EV-mediated delivery of the murine immunomodulatory molecule Ox40L shows significant therapeutic efficacy in a tumor model *in vivo*

After the successful establishment of a functional EV bioengineering platform for mRNA loading and delivery using CD63-PUFe and co-display of VSVg, we next adapted the system for the delivery of a therapeutically relevant mRNA *in vivo*.

One of the most promising strategies for sustainable cancer eradication is immunotherapy, including the induction of the OX40/OX40L axis, a co-stimulatory immune cell molecule interaction potentially creating cancer-specific memory T cells for prolonged anti-tumor responses^48–51^. Therefore, we hypothesized that the injection of EVs actively loaded with murine Ox40L (mOx40L) mRNA into tumors of the highly aggressive B16F10 melanoma model would elicit a sustainable mOx40L display on tumor cells. Moreover, the co-delivery of mRNA and passively loaded protein was expected to increase the therapeutic response due to its dual short and long-term mode of action (Figure 5a and 5b).

**Figure 5.**
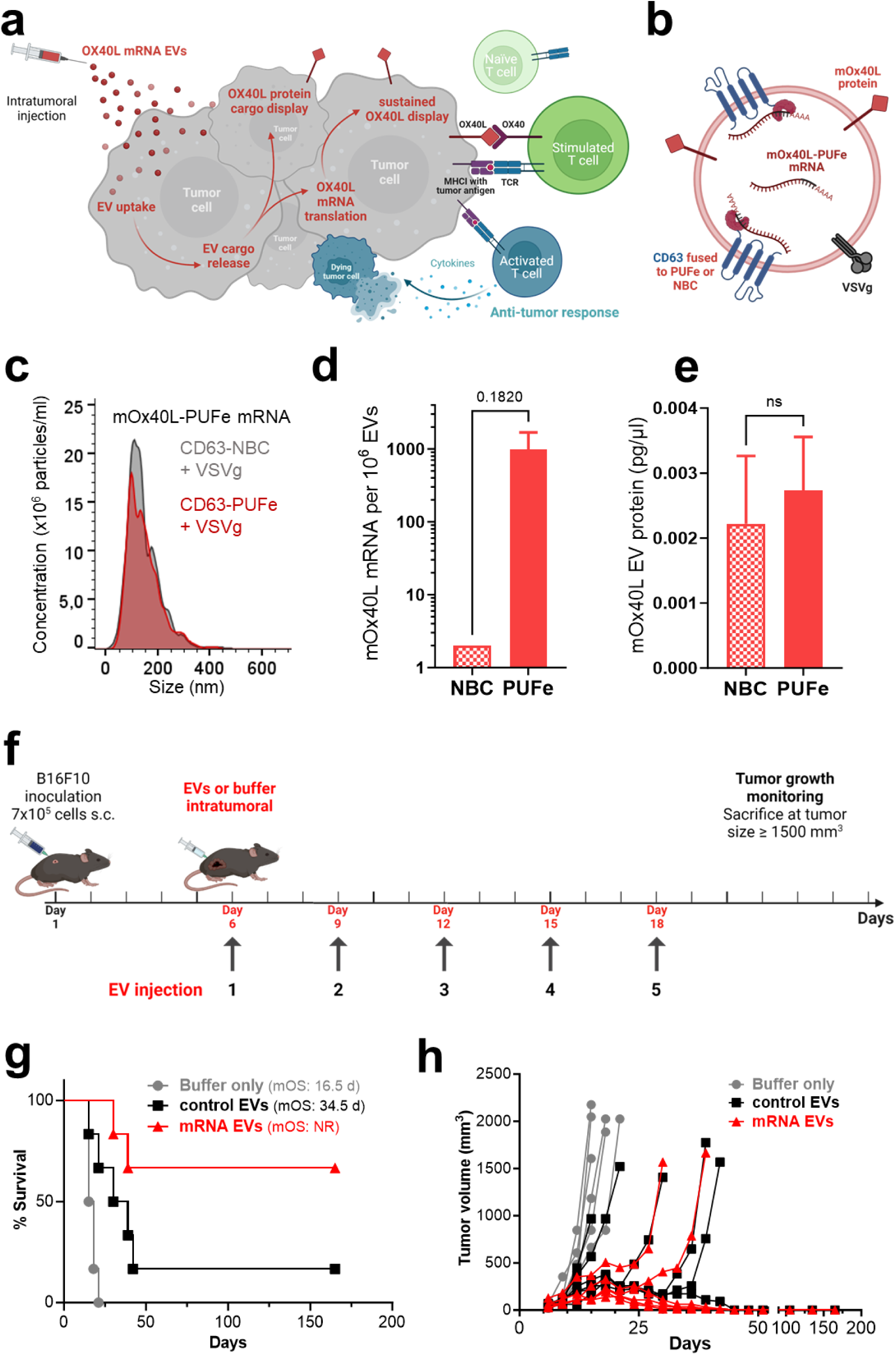
Efficient EV-mediated delivery of the immunomodulatory molecule mOx40L showed significant therapeutic impact in murine tumor model *in vivo*. (a) Illustration of the proposed therapeutic short-term and long-term dual effect of EV-mediated delivery of mOx40L protein and mRNA. Co-loaded mOx40L protein is displayed immediately at the surface of targeted tumor cells, while the delivered mRNA is translated into protein for a sustained co-stimulatory immunotherapeutic effect leading to an anti-tumor response. Figure created using BioRender. (b) Cargo composition of engineered mOx40L mRNA and protein-loaded EVs. (c) Average particle size determination by NTA of mOx40L-PUFe mRNA EVs and control EVs with VSVg co-expression. (d) Absolute quantification of mOx40L-PUFe mRNA molecules per 10^6^ EVs. (e) Concentration of mOx40L protein in mOx40L-PUFe mRNA and control EVs as measured by murine Ox40L sandwich ELISA. (f) Injection scheme for intratumoral injections of mRNA EVs, control EVs, or suspension buffer (PBS-HAT) at a dose of 2 ng mRNA per kg bodyweight into B16F10 melanoma-bearing mice (n=6 per group). After tumor engraftment, mice were injected 5 times and monitored regularly for tumor growth. Figure created using BioRender. (g) Kaplan-Meier survival analysis of mice treated with mOx40L mRNA EVs, control EVs, or buffer only with assessment of median overall survival (mOS). All curves are significantly different from each other (Log-rank (Mantel-Cox) test, P value 0.0003. NR – not registered. (h) Tumor volumes measured regularly after the last injection. Each line represents one mouse of the respective group. Four out of six of the mRNA EV-treated mice and one out of six of the control EV treated mice went into complete remission and lost their tumor beyond palpability for the duration of the experiment (165 days). Curve analysis (Wilcoxon Signed Rank Test): Buffer only group P value (two-tailed) 0.0312, Control EV group P value (two-tailed) 0.0002, mRNA EV group P value (two-tailed) 0.0002, α=0.05, all curves significant.

Control EVs and mOx40L-PUFe mRNA EVs displaying VSVg were produced from cells stably expressing the mRNA and mean particle size was validated at 121.3 ± 7.2 nm (Figure 5c), and presence of EVs was verified by electron microscopy (Supplementary Figure S 3a). Active EV loading of mOx40L mRNA was confirmed by RT-qPCR (Figure 5d). Next, the co-loading of mOx40L protein in both mOx40L-PUFe mRNA-loaded and control EVs was measured by ELISA (Figure 5e) and EV flow cytometry (Figure S 4b – S 4e), respectively. For both mRNA-loaded and control EVs the same amount of passive mOX40L protein loading was observed (Figure 5e and Supplementary Figure S 4d) and the detected mOx40L protein was associated with the EV membrane (Supplementary Figure S 4b and S 4c).

To test our hypothesis *in vivo* mRNA EVs at a dose of 2 ng mOx40L mRNA per kg bodyweight, particle count-matched control EVs, or buffer were injected intratumorally into mice with engrafted B16F10 melanoma. At first, three injections were applied every third day after tumor engraftment (n=5 animals per group) as illustrated in Supplementary Figure S 5a. The experiment was terminated 30 days after the first injection. All mice of the buffer-only group (median overall survival 18 days) and control EV group (median overall survival 23 days) were sacrificed due to reaching the humane endpoint (Supplementary Figure S 5b and S 5c). Mice injected with mRNA-loaded EVs showed a significantly slower tumor growth than control EV-injected or buffer-injected mice (Supplementary Figure S 5c). Importantly, two mice (40%) of the mRNA EV group, went into clinical complete remission and lost their tumors at the time of experiment termination (Supplementary Figure S 5b and S 5c). These data prompted us to perform a subsequent experiment with longer exposure to treatment (5 injections, Figure 5f). Here, non-treated mice (buffer-only group) showed a similarly aggressive rate of tumor growth as before and a median overall survival of 16.5 days (Figure 5g). Control EV-injected mice, however, displayed a slower tumor growth and a longer median overall survival than in the previous experiment, 34.5 days, while the mRNA treatment group outperformed both control groups (Figure 5g and 5h). Intriguingly, one mouse (17%) of the control EV group and four mice (67%) of the mRNA EV group went into complete remission 45 days after the first injection and lost their tumors beyond palpability (Figure 5h). These 5 surviving mice were monitored regularly for relapse of tumor growth, but none was detected until the experiment was terminated 165 days after the initial injection (Figure 5g and 5h).

These results impressively showed the therapeutic impact of mOx40L expression on tumor cells *in vivo* and suggest a high versatility of the EV mRNA loading platform as a therapeutic approach for cancer immunotherapy.

## DISCUSSION

Functional delivery of therapeutic mRNA faces major challenges, including unfavorable drug stability, immunogenicity, and inefficient delivery, leading to insufficient therapeutic mRNA abundance at the target site. The data presented here address these matters by combining the use of EVs as nature’s very own nanoparticles with state-of-the-art synthetic biology strategies to develop a highly promising new platform for functional mRNA delivery. To maximize Nanoluc mRNA loading into EVs, a highly efficient EV sorting platform consisting of the EV-enriched tetraspanin CD63 fused to an engineered version of the high-affinity RNA-binding protein (RBD) PUF^37^, termed PUFe, was designed. We showed that CD63-PUFe efficiently loaded mRNA with the compatible PUFe-binding sequence in its 3’UTR into the lumen of engineered EVs. The observed mRNA delivery *in vitro*, however, did not reflect the significant enrichment of Nanoluc-PUFe mRNA EV cargo. Another major obstacle for functional delivery of mRNA therapeutics is inefficient endosomal escape in the target cell. Thus, VSVg, a viral protein naturally capable of fusing the viral envelope with endosomal membranes and used to enhance EV cargo delivery previously^33,46^, was incorporated into our EV engineering strategy. With this modification, Nanoluc mRNA delivery was substantially improved *in vitro* and *in vivo*. Next, functional mRNA cargo delivery was clearly proven by refining the engineering strategy for Cre recombinase mRNA, showcasing the functionality of our EV-based mRNA loading platform for gene editing modalities. Finally, the therapeutic impact of EVs loaded with the murine immunostimulatory co-receptor Ox40L mRNA and protein was demonstrated in the highly aggressive B16F10 melanoma model. When mRNA-loaded EVs were injected five times intratumorally, 4 out of 6 mice had no palpable tumor left at 45 days post injection and survived 165 days with no signs of tumor relapse when the experiment was terminated.

As demonstrated by the passive loading control throughout the study, mere overexpression of the target mRNA is not sufficient for effective endogenous mRNA loading into EVs. Instead, we and others implemented active endogenous loading strategies to enrich target mRNA in EVs^28,29,33,43^. However, most studies use producer cell engineering strategies based on transient co-transfection of the target mRNA-encoding plasmid. This approach leads to the potential co-isolation of transfection complexes containing the mRNA-encoding plasmid with the EVs and could thus cause misinterpretation of experimental results from functional studies^34–36^. Our strategy, on the other hand, takes advantage of the non-viral stable genomic integration of the target mRNA expression cassette using the ϕC31 integrase system^44, 45^, and thereby ensures that the target mRNA can only be delivered by EVs, and not residual transfection complexes. Consequently, our platform includes passively loaded control EVs produced under equal biological conditions from the same producer cells and crucial for the setting of accurate baselines for the co-delivery of passively loaded mRNA and protein.

To address a substantial challenge in therapeutic EV cargo delivery, efficient endosomal escape, VSVg was incorporated into the membrane of CD63-PUFe or control engineered EVs. However, there are risks associated with the use of VSVg, mostly related to safety concerns due to its viral origin, and thus the implementation of alternative fusogenic proteins would be advantageous before further advancing this system towards clinical application.

Interestingly, the data presented in this study revealed a major incongruence between the enrichment of target mRNA cargo in engineered EVs and the observed cargo delivery efficiencies *in vitro* and *in vivo*, even with additional engineering of VSVg co-display for enhanced endosomal escape. In contrast, locally high concentrations and prolonged dosing of therapeutic EVs by repeated intratumoral injections of EVs returned a highly favorable cancer immunotherapeutic outcome in an aggressive mouse tumor model. For systemic application, however, the absolute mRNA molecule numbers per EV might still be too low for exerting relevant therapeutic effects in a clinical setting. Thus, further improvements of the platform could be implemented by fusing other highly abundant EV proteins to PUFe^22^, employing combinatorial approaches to engineer a multitude of EV subpopulations^22^, introducing stabilizing moieties^21^, and displaying targeting antibodies for directed therapeutic EV cargo delivery^23^.

The current state-of-the-art delivery vehicles for nucleic acid therapeutics are synthetic lipid-based nanoparticles, such as LNPs. However, one major drawback of the use of synthetic LNPs is the inefficient delivery into extrahepatic tissues. Here, we showed EV-mediated extrahepatic mRNA delivery at a dose at least one order of magnitude lower than currently used for LNP-mediated mRNA delivery^52^. In fact, superior RNA delivery efficiencies *in vivo* by EVs as compared to LNPs has been reported previously using different EV engineering strategies^24,53, 54^. Here, we successfully treated mice implanted with highly aggressive B16F10 melanoma with EVs loaded with effective cancer immunotherapy, the murine immunostimulatory co-receptor Ox40L mRNA and protein. To our knowledge, this was the first time such a high response rate was achieved with durable complete remission in the B16F10 melanoma model. In comparison, intratumorally injected LNPs delivering OX40L mRNA in combination with IL-23 and IL-36γ mRNA in the same melanoma model resulted in only 40% of the mice being alive after 35 days, despite excellent delivery of the mRNA^55^. Importantly, these LNPs were formulated at a dose of 1.67 µg OX40L mRNA per injection^55^, which is more than 4 orders of magnitude higher than the amount of Ox40L mRNA (58 pg per injection, corresponding to 2 ng per kg bodyweight) delivered by EVs produced with our technology. This strongly support the notion that EVs are much more efficient in nucleic acid therapeutics delivery than LNPs, and that the dual short- and long-term effect of co-delivered protein with mRNA-loaded EVs is highly beneficial for the overall therapeutic outcome. Therefore, the results presented in this study are an indispensable addition to the drug delivery field demonstrating the exceptional versatility of EVs engineered with our loading platform as functional mRNA delivery vehicles.

## METHODS

### Cell culture

HEK293T, Huh7, T47D-Traffic Light, B16F10-Traffic Light cells were cultured in DMEM medium with high glucose, GlutaMAX and pyruvate (Gibco). The medium was additionally supplemented with 1% Anti-Anti solution (Gibco) and 10% fetal bovine serum (FBS, Gibco). Cells were cultured in a humidified atmosphere at 37℃ and 5% CO_2_. All cells were kept at low passages (< 15 passages) and high viability (> 90%).

### Plasmids and construct generation

All overexpression constructs (EV scaffold and mRNA) were codon-optimized to ensure optimal expression and bought from Thermo Fisher Scientific already cloned into their Thermo vector backbone. All constructs were subcloned into the pLEX vector backbone using EcoRI and NotI restriction enzymes for transfection-based EV production. For the generation of mRNA stable cells, mRNA constructs were transferred to the FC550A-1 backbone (System Biosciences) belonging to their ϕC31 integrase vector system using the same restriction sites. VSVg was expressed from the pMD2.G plasmid (Addgene #12259).

For the comparison to the SEND platform, the published expression vectors (pCMV-HsPeg10rc3, #174859; pCMV-Hs.cargo(Cre), #174863) were purchased from Addgene and their sequences were validated by restriction enzyme digestion and partial re-sequencing.

### Transient transfection

For large-scale transfections and EV production, HEK293T cells were seeded in 150-mm cell culture dishes or 10-layer cell factories one day prior to reach a confluency of 70% at the time of transfection. Cells were either transfected using PEI Max 40K (Polysciences) at a DNA:PEI ratio of 1:3 (w/w) or Lipofectamine 2000 (Thermo Fisher Scientific) at a DNA:Lipofectamine ratio of 1:2 (w/v). Transfection complexes were allowed to form in serum-free defined medium (OptiMEM, Gibco), and added dropwise to the cells. After 4 h, transfection medium was removed and changed to OptiMEM as EV production medium.

For small-scale plasmid transfections, cells were seeded at an appropriate amount in a multi-well plate one day prior to the time of transfection. When the cells reached a 70% confluency, transfection medium containing plasmid DNA and Lipofectamine 2000 at a ratio of 1:2 (w/v) was added to the culture medium. The cells were harvested at 24 h post-transfection for expression analysis.

For EV production for the SEND platform comparison, HEK293T cells were transfected with PEI Max 40K (Polysciences) and medium was either changed to OptiMEM, as described above, or changed to fresh full growth medium as described in Segel et al., 2021, supplementary material^30^.

### ϕC31 integrase stable cell generation

HEK293T cells stably expressing mRNA constructs were generated using the ϕC31 integrase system (System Biosciences) according to the manufacturers protocol. Briefly, HEK293T cells were co-transfected with the donor vector FC550A-1 containing the mRNA of interest flanked by attB donor sites and the integrase vector FC200PA-1 encoding for the ϕC31 integrase enzyme using Lipofectamine 2000 Transfection reagent at a ratio of 1:4 (w/v DNA:Lipofectamine 2000). The transient expression of the enzyme leads to a single-copy non-viral transgene integration into the host cell genome by site-specific recombination of the attB sites from the donor plasmid with a pseudo attP site in the genome^40, 41^. The integrated cassette also encodes for the puromycin resistance gene. 48 h after transfection, Puromycin (1 µg/ml) was added to the culture medium to select for successfully transfected cells. Selection was completed at 2 weeks post-transfection, and cells were analyzed for genomic vector integration and transgene expression. For culture, stable cells were kept in full growth medium constantly supplemented with Puromycin at a final concentration of 4 µg/ml.

### EV production

Depending on the application, EVs were produced from cells cultured either in 150-mm cell culture dishes (12×10^6^ cells/plate, 20 ml culture medium) or 10-layer cell factories (4.5×10^8^ cells/factory, 600 ml culture medium). After transient transfection, all medium was removed and HEK293T cells were cultivated in serum-free OptiMEM (Gibco). After 48 h, conditioned medium was collected and subjected to one of our streamlined differential centrifugation and ultrafiltration protocols for EV isolation. Briefly, medium was spun at 700xg for 5 min to remove residual cells, then at 2,000 xg for 10 min to remove cell debris. The centrifuged medium was then filtered through a polyethersulfone (PES) membrane filter with 0.22 µm pore size (TPP). The following concentration steps were chosen according to experimental conditions and medium volumes. For small scale volumes and many samples, filtered conditioned media were subjected to ultracentrifugation at 100,000 xg for 90 min at 4℃ in a Beckman Coulter Optima XP Ultracentrifuge. For large scale volumes and few samples, the conditioned medium was run through a hollow-fibre filter (D06-E300-05-N, MIDIKROS 65CM 300 K MPES 0.5 MM, Spectrum Laboratories) using a tangential flow filtration system (KR2i TFF System, Spectrum Laboratories) at a flow rate of 100 ml/min with a transmembrane pressure at 3.0 psi and shear rate at 3700/sec. After diafiltration with PBS, the eluate was reduced to approximately 40–50 ml and then concentrated using an Amicon Ultra-15 10 kDa molecular mass cut-off spin filter (Merck Millipore). Final EV preparations were kept in sterile filtered PBS for immediate use or sterile PBS-HAT buffer^56^ for storage at −80℃.

### Nanoparticle Tracking Analysis

Nanoparticle tracking analysis (NTA)^57, 58^ is based on the motion of sub micrometer-sized particles in solution (Brownian motion) and was used for quantifying the concentration and size distribution of particles in all EV samples. NTA was performed using the NS500 instrument equipped with the NTA 2.3 analytical software (NanoSight). After isolation, an aliquot of each EV sample was diluted in 0.22 μm filtered PBS between 1:500 to 1:5000 to achieve an appropriate particle count of between 2×10^8^ and 2×10^9^ per ml for the measurement. The camera focus was adjusted to make the particles appear as sharp dots. Using the script control function, five 30 s videos for each sample were recorded, incorporating a sample advance and a 5 sec delay between each recording. All recordings were done in light scatter mode at a camera level of 13-14. The analysis of all EV measurements was performed with constant software settings, screen gain at 10 and detection threshold at 7.

### RNA isolation

A defined amount of EVs, such as 1×10^10^, or 3-5×10^5^ pelleted cells were lysed in TRI Reagent (“LS” for fluid EV samples, conventional for cells, Merck Millipore). Liquid chloroform was added to the lysates at a 1:5 v/v ratio and vigorously mixed for 30 s. Lysates were incubated at room temperature for 3-5 min to initiate phase separation and centrifuged at 4℃ for 15 min at 12,000xg. The RNA-containing aqueous phase was carefully collected without disturbance of the DNA-containing organic phase and transferred to a fresh microtube. The RNA was mixed with ethanol absolute at a 1:1 v/v ratio and submitted to a spin column-based purification protocol (Direct-zol RNA Miniprep kit, Zymo Research), including a DNAse treatment step, according to the manufacturer’s instructions. RNA concentration was measured using the Implen NanoPhotometer NP80 (Implen).

### RT-qPCR

A defined amount of RNA (for EV RNA at least 40 ng, for cellular RNA at least 100 ng) was subjected to cDNA synthesis using the High-Capacity cDNA Reverse Transcription Kit (Thermo Fisher Scientific) and a 15-mer oligo-dT primer according to the manufacturer’s protocol. Briefly, 10 µl of a 2X RT master mix consisting of ddH_2_O, 2X RT buffer, 8 mM dNTPs, 20 µM oligo-dT primer and 50 U MultiScribe Reverse Transcriptase was mixed with 10 µl RNA and incubated for 10 min at 25℃ followed by 2 h at 37℃ in a thermocycler. The reaction was stopped by heating to 85℃ for 5 min. Depending on the initial RNA input, cDNA samples were diluted with ddH_2_O so that 1 µl cDNA corresponded to 2 ng RNA for EVs or 5 ng RNA for cells. Quantitative real-time PCR (qPCR) was performed using the PowerUp SYBR Green Master Mix (Thermo Fisher Scientific), a pre-formulated universal 2X master mix containing Dual-Lock Taq DNA polymerase for real-time PCR. Briefly, a 10 µl qPCR reaction was set up using 5 µl 2X master mix, forward primer at 200 nM final concentration, reverse primer at 200 nM final concentration, 1 µl cDNA and ddH_2_O. Samples and standards were measured in triplicates using the CFX96 Touch Real-Time PCR Detection System (BioRad). Cycling conditions were used as follows: 1. 50℃ for 2 min, 2. 95℃ for 2 min, 3. 95℃ for 15 s, 4. 62℃ for 1 min + plate read, 5. go to 3 for 39 more times, 6. 95℃ for 15 s, 7. melt curve from 60℃ to 95℃ (increment 0.5℃) for 5 s + plate read. Primers were designed to amplify the 5’end of the respective mRNA, thus only full-length transcripts were quantified. The primer sequences are listed in Table 1:

**Table 1:**
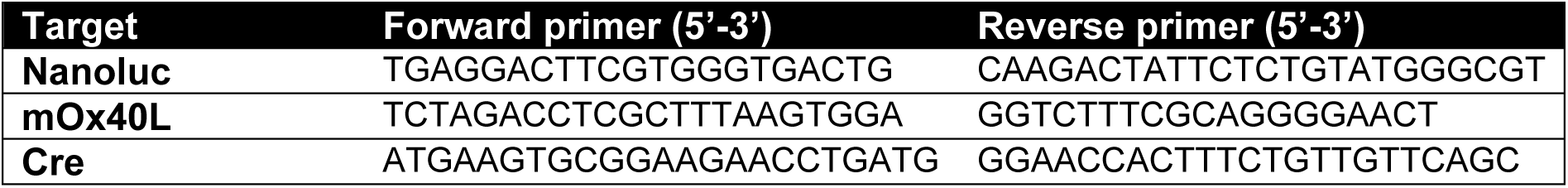
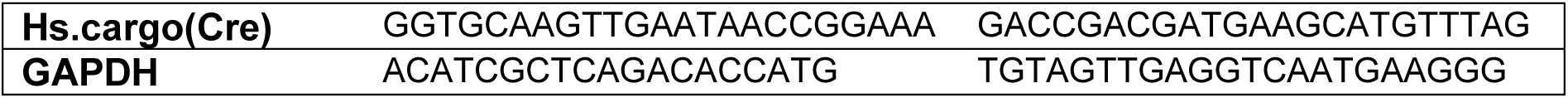
Gene-specific primer sequences for mRNA quantification by RT-qPCR.

### Relative and absolute quantification of RT-qPCR data

Full-length target mRNA in producer cells were measured using the 2^-ΔΔCq^ relative quantification method with GAPDH mRNA expression as reference.

As there is no validated reference mRNA for EV mRNA analysis established yet, applied absolute quantification of Nanoluc mRNA molecules in RNA from EVs was performed using standard curves with known molecular concentrations generated for each primer pair. The target product was PCR-amplified from an appropriate template, such as the expression plasmid or cDNA, using the HotStarTaq Plus Master Mix Kit (Qiagen) as follows: 1. 95℃ for 5 min, 2. 94℃ for 30 s, 3. Primer-defined annealing temperature for 30 s, 4. 72℃ for 1 min, 5. go to 2 for 30 more times, 6. 72℃ for 10 min. The PCR product was run on a 1% agarose gel and subsequently purified using the NucleoSpin Gel and PCR Cleanup kit (Macherey-Nagel) according to the manufacturer’s protocol. The DNA concentration of the purified PCR product was measured using the NanoPhotometer NP80 (Implen) and diluted in a solution of murine RNA at 2 ng/µl in a series of 10X-steps up to 10^-9^-fold. The optimal range of standard dilution series was determined for each primer pair and usually observed between the 1:10^-4^ and 1:10^-7^ dilutions. For each assay of qPCR performed on sample cDNA derived from EV RNA or cellular RNA, at least 3 serial dilutions of the respective standard were analyzed on the same plate. The Cq values of the standards were correlated to their absolute molecule count per reaction as determined by the initial template concentration, amplicon length and molecular weight, and final dilution. For every primer pair a standard curve was calculated using linear regression. Then, the standard curve equation was applied to the samples. Apart from the Cq values measured and their quantification according to the standard curve, the calculations of absolute mRNA molecules in a sample included the total amount of isolated RNA, the RNA amount used in reverse transcription as well as the RNA amount corresponding to the cDNA per qPCR reaction. Lastly, the absolute molecules were normalized per cell or per 1×10^6^ EVs.

### RNA transfection

Huh7 or HEK293T recipient cells were seeded at 1×10^5^ cells per well in a 24-well plate. The following day, 125 ng RNA purified from either EVs or corresponding producer cells were transfected per well in serum-free OptiMEM (Gibco) using Lipofectamine 2000 at a ratio of 1:2 (w/v), i.e. 0.25 µl per transfection. After 30 min incubation at room temperature, transfection complexes were added directly to the culture medium. Cells were harvested and lysed for protein measurement and Nanoglo luciferase assay at 3, 6, 12, 24, and 48 h post-transfection.

### NanoGlo luciferase assay

The NanoGlo luciferase assay kit (Promega) was used to determine the amount of functional Nanoluc protein in lysed biological samples by supplying the enzyme with substrate and immediately measuring the resulting luminescence. Cells were lysed directly in the well of the cell culture plate in an appropriate amount of PBS supplemented with 0.1% Triton-×100. Lysis was performed on ice for 30 minutes. Lysates were collected and cleared by centrifugation. The protein concentration was measured using the DC protein assay kit (BioRad) according to the manufacturers protocol.

Tissue samples from internal organs were processed as previously described^19^. In brief, organs were isolated, weighed, and suspended in 1 ml ice-cold PBS supplemented with 0.1% Triton-×100. Mechanical disruption was performed using the TissueLyser II and 8-mm stainless steel beads (both Qiagen) at an amplitude of 30 for 2 min at a time. After each lysis run, samples were rested on ice and viscosity was tested by pipetting up and down. Lysis steps were repeated as necessary until lysates were homogenous without obvious remains of solid tissue.

25 µl EVs, cell lysate, or diluted tissue lysate from murine internal organs were used for Nanoluc luciferase detection. The EVs or lysates were plated in opaque 96-well microplates. NanoGlo reagent (NanoGlo Luciferase Assay System, Promega) was prepared from NanoGlo lysis buffer and NanoGlo substrate solution according to the manufacturer’s instructions. Using the GloMax 96 Microplate Luminometer (Promega), 25 μl NanoGlo reagent were added to each well by automatic injection and immediately analyzed.

### Western Blot

For Western Blot, cellular protein was extracted from 3-5×10^5^ pelleted cells. The cells were lysed on ice for 30 min in 100 – 200 µl RIPA buffer supplemented with 1X cOmplete protease inhibitor cocktail (Roche). EVs were directly lysed in PBS/0.1% Triton-×100. The protein amount of the cleared lysates was measured using the DC protein assay kit (BioRad) according to the manufacturers protocol. Prior to SDS-PAGE, all samples were heated for 5 min at 70℃ in 1X SDS-PAGE loading buffer (0.4 M sodium carbonate, 10% glycerol, 0.5 M dithiothreitol, and 8% SDS). Then, NuPAGE Novex 4–12% Bis-Tris Protein Gels (Thermo Fisher Scientific) were loaded with denatured samples, a molecular size marker (PageRuler Plus prestained, Thermo Fisher Scientific), and run in NuPAGE MES SDS running buffer (Thermo Fisher Scientific) at 120 V for at least 2 h. Semi-dry blotting was performed using the pre-stacked iBlot system (Thermo Fisher Scientific) to transfer the proteins to a pre-activated nitrocellulose membrane. The membranes were blocked with Intercept (PBS) Blocking Buffer (LI-COR Biosciences) for 50 min at RT. Primary antibodies were added at appropriate concentrations to the membranes and incubated at 4℃ for 16-24 h on a horizontal shaker. After primary antibody removal, the membranes were washed three times for 10 min in PBS-T (PBS with 0.1% Tween-20, Merck). Then, the appropriate IRDye secondary antibodies (LI-COR Biosciences) were added to the membranes and incubated for 50 min at RT. Next, the membranes were washed three times for 10 min in PBS-T and visualized by scanning both 700 nm and 800 nm channels on the LI-COR Odyssey CLx infrared imaging system (LI-COR Biosciences).

The primary antibodies used in this study were the following: rb-α-Nanoluc (non-commercial, Promega, 1:1000); rb-α-CD63 (ab68418, Abcam, 1:1000), gt-α-VSV-G Tag (PA1-30278,

Thermo Fisher Scientific, 1:1000), rb-α-Cre recombinase (ab188568, Abcam, 1:1000), rb-α-PEG10 (#77111, Cell Signaling Technology, 1:1000), ms-α-SDCBP (Syntenin) (TA504796, Thermo Fisher Scientific, 1:500), ms-α-ACTB (A5441, Merck, 1:20000); rb-α-Calnexin (ab22595, Abcam, 1:1000).

The secondary antibodies used here were: gt-α-ms IRDye800CW or 680LT (1:10000), gt-α-rb IRDye800CW or 680LT (1:10000), and dy-α-gt IRDye680RD (1:10000) (all LI-COR Biosciences).

### EV uptake *in vitro*

For quantification of cellular uptake of Nanoluc-PUFe mRNA-loaded EVs *in vitro*, human hepatocellular carcinoma (Huh7) cells were seeded at a density of 6×10^4^ cells per well in a 24-well plate. The following day, the culture medium was renewed and EVs were added directly to the culture medium. Per test, equal amounts of EVs (mRNA-loaded or control), ranging from 1×10^8^ to 7.5×10^10^ per well depending on loading efficiencies and scope of the experiment, were used to eventually assess the excess Nanoluc protein signal of mRNA translation over pre-existent passively loaded protein. EVs were co-cultured with the cells for 2 to 48 h, as indicated per experiment. Then, the EV-containing medium was removed, the cells were washed extensively in PBS (at least 3 times for 15 minutes each) by gentle horizontal shaking, followed by direct lysis on the plate with PBS-TritonX 100 (0.1%). Then, cell lysates and conditioned media were subjected to the NanoGlo luciferase assay as described. For every treatment an “EV only”-control well was included: a cell culture well with growth medium, but without cells, was equally treated with the corresponding amount of EVs and the signal from the medium served as normalization control to account for the total protein delivered by the EVs without translation.

For quantification of cellular uptake and activity of Cre mRNA-loaded EVs *in vitro*, human T47D-Traffic Light and murine B16F10-Traffic Light cells were seeded in a 96-well plate at a density of 1×10^4^ cells per well. The following day, the cell culture medium was renewed and equal amounts of mRNA-loaded and control-loaded EVs, ranging from 1×10^8^ to 1×10^10^ EVs per well depending on loading efficiencies and scope of the experiment, were added to the cells. EVs were co-cultured with cells for 24 to 72 h, as indicated per experiment. For analysis, EV-containing medium was removed, cells were washed once with PBS and then enzymatically detached with 0.05% Trypsin (Gibco) to be measured by flow cytometry for the frequency and intensity of eGFP expression.

### SEND platform comparison

For the SEND platform comparison experiments, EVs were produced either according to our streamlined workflow (see above) or as described in Segel et al., 2021^30^, supplementary material. In brief, HEK293T cells were seeded in 20 ml of full growth medium in 150-mm dishes at a density of 12×10^6^ cells/plate to reach 70% confluency at the time of transfection. For production of SEND(bind) or SEND(control) EVs, cells were transfected with a total of 45 µg plasmid DNA per plate consisting of 15 µg pCMV-Hs.cargo(Cre), 15 µg pCMV-hs.PEG10rc3 (SEND(bind)) or empty vector backbone (SEND(control)), and 15 µg pMD2.G (VSVg expression). Accordingly, to produce Cre-PUFe (bind) and Cre-PUFe (control) EVs cells were transfected with 15 µg pLEX-Cre-PUFe_bind_, 15 µg pLEX-CD63-PUFe (bind) or pLEX-CD63-MCP (control), and 15 µg pMD2.G (VSVg expression). Four hours after transfection, medium was changed either to OptiMEM (our protocol) or replaced with fresh full growth medium (Segel et al. protocol^30^) and cells were grown for 48 hours. Next, OptiMEM conditioned media were subjected to our protocol (see above), or as follows (Segel et al. protocol): low centrifugation at 2000 xg for 10 minutes, filtration through a polyethersulfone (PES) membrane filter with 0.45 µm pore size (VWR), Ultracentrifugation at 4°C and 120,000 xg for 2 hours, supernatant decanting, and resuspension in 200 µl sterile PBS per harvested plate.

Particle sizes and concentrations were measured by NTA for EVs produced with our protocol, as the input conditioned medium was free from supplements and serum. Uptake experiments in Traffic Light Cre reporter cells were performed as described.

As the input conditioned medium for EVs produced by the Segel et al. protocol was supplemented with FBS, no NTA measurements were performed due to light scattering inaccuracies caused by large serum protein complexes. Instead, uptake experiments were performed with equal volumes of EV suspension after ultracentrifugation. Here, 1×10^4^ Traffic Light Cre reporter cells in a 96-well plate format were incubated with 20 µl EV suspension for 72 hours, as published by Segel et al.^30^.

### Virus production

The pLV-CMV-LoxP-DsRed-LoxP-eGFP plasmid used to produce virus (termed Traffic Light virus) was purchased from Addgene (#65726). HEK-293T cells were seeded into T175 flask at the density of 20 million/ flask. pCD/NL-BH and pcoPE01 as the helper and envelope plasmid respectively, were co-transfected into the HEK-293T cells on the second day. The amount of plasmids used per flask was 22.5 µg for pLV-CMV-LoxP-DsRed-LoxP-eGFP and pCD/NL-BH, and 7 µg for pcoPE01. After co-transfection overnight, the medium was changed to full DMEM (10% FBS and 1% Anti-anti) with 10mM sodium butyrate (Sigma-Aldrich) to boost CMV promoter activity. After boosting for 8 hours, the medium was changed back to full DMEM without sodium butyrate. And then the viruses were harvested 24 hours after the second time medium change. Briefly, the viruses-containing medium was filtered through a 0.45 µm syringe filter (VWR) to the Nalgene® Oak Ridge Centrifuge Tubes (Thermo Scientific) before centrifuging at 25,000x g for 90 min at 4°C. Afterwards, the supernatant was discarded, and the virus pellets were resuspended by 1ml freshly prepared medium (IMDM with 20%FBS) before storing them at −80°C freezer for long-term use.

### Traffic Light stable cell generation

B16F10 and T47D cells were seeded into 6-well plates 1 day before adding the Traffic Light virus. 2 doses (10 µl and 50 µl) of viruses were added into the candidate cell lines. After 2 days of transduction, 2 µg/ml and 4 µg/ml puromycin were used to select the positively transduced B16F10 and T47D cells respectively. Around 1 week later, the survived cells formed colonies and the untransduced cells died from puromycin pressure. If the cells survived for the high dose (50 µl) of virus, use these cells for further selection. Otherwise, use the cells transduced with low dose (10 µl) of virus. The colonies were trypsinized and transferred to T25 flasks and cultured with puromycin pressure continuously. After 2 passages of the selection, the stable cells were ready for downstream use.

### Flow cytometry

For flow cytometry analysis, adherent cells were enzymatically detached from the culture dish using 0.05% Trypsin (Gibco) and collected in microtubes using an appropriate amount of full growth medium. The cells were spun down at 900xg for 5 min and resuspended in 100 μL of full growth medium or PBS supplemented with 2% FBS. If applicable, cells were stained with the appropriate amount of fluorescent antibody for 30 min in the dark at 4℃, washed with 1 ml PBS, spun down at 900xg for 5 min, and resuspended in 100 μL of full growth medium or PBS supplemented with 2% FBS. Cells were transferred to a 96-well V-bottom microplate (Sarstedt) for flow cytometry analysis using a MACSQuant Analyzer 10 instrument (Miltenyi Biotec). Dead cells were excluded from analysis via 4,6-diamidino-2-phenylindole (DAPI) staining and doublets were excluded by forward/side scatter area versus height gating. Analysis was performed using FlowJo Analysis Platform (BD Biosciences), versions 10.7 and 10.8.

Flow cytometry antibodies used in this study: Invitrogen anti-mouse OX40L Monoclonal Antibody (OX89)-FITC, Thermo Fisher Scientific.

### High Resolution Single EV analysis by Imaging Flow Cytometry

For single EV analysis experiments by Imaging Flow Cytometry (IFCM), EV samples were diluted in PBS-HAT (DBPS supplemented with 25 µM HEPES, 0.2% human albumin and 25 µM trehalose)^56^ to a final concentration of 1×10^10^ particles/mL before usage. A volume of 25 µL (equivalent to 2.5 x 10^8^ particles) was incubated with anti-mouse OX40L-FITC antibodies (Invitrogen anti-mouse OX40L Monoclonal Antibody (OX89)-FITC, Thermo Fisher Scientific, #MA-5-17912) at a final antibody concentration of 4 nM overnight. Post staining, samples were diluted 1:2,000 in PBS-HAT before acquisition on a Cellstream instrument (Amnis/Cytek) with FSC turned off, SSC laser set to 40%, and all other lasers set to 100% of the maximum power. Small EVs were defined as SSC(low) by using CD63-mNeonGreen (mNG)-tagged EVs as biological reference material as described before^59^, and regions to quantify fluorescence-positive populations were set according to unstained samples. Samples were acquired for 5 minutes at a flow rate of 3.66 µL/min (setting: slow) with CellStream software version 1.2.3 and analyzed with FlowJo Software version 10.5.3 (FlowJo, LLC). Dulbecco’s PBS pH 7.4 (Gibco) was used as sheath fluid. Fluorescence calibration was performed as described previously.^56,59, 60^ In brief, FITC MESF beads (Quantum FITC-5 MESF, Bangs Laboratories Inc., cat 555A, lot 13734) and APC MESF beads (Quantum APC MESF, Bangs Laboratories Inc., cat 823A, lot 13691) with known absolute fluorescence values for each bead population were acquired with the same settings used for EV measurements with the exception that the SSC laser was turned off, and linear regressions were performed to convert fluorescence values into FITC/APCMESF values, respectively. Flow cytometric plots using MESF unit axes were created with FlowJo v 10.5.3 (FlowJo, LLC).

### mOx40L ELISA

To measure the mOx40L protein abundance in EVs the Mouse TNFSF4 solid-phase sandwich ELISA (enzyme-linked immunosorbent assay) kit (Invitrogen, Thermo Fisher Scientific) was used according to the manufacturer’s instructions. In brief, 100 µl of standard and 100 µl of appropriately diluted sample were bound to the solid phase in capture antibody-coated microtiter plates. The EV samples were measured in several dilution steps, ranging from 100-fold to 10,000-fold dilutions. After washing, the detector antibody (biotinylated conjugate) was bound to the antigen for sandwich formation. Next, the biotinylated antibody-antigen sandwich was linked to the HRP (Horse radish peroxidase) enzyme via Streptavidin and then the chromogenic substrate 3,3′,5,5′-Tetramethylbenzidine (TMB) was supplied for detection. Subsequently, the enzymatic reaction was stopped and the absorbance at 450 nm was measured with a CLARIOstar Plus microplate reader (BMG Labtech). The intensity of this signal is directly proportional to the concentration of murine Ox40L target protein present in the original specimen. Murine Ox4L target protein concentration was determined by extrapolation from a standard curve obtained by linear regression of the standard measurement values.

### Negative stain transmission electron microscopy (nsTEM)

Three microliters of the sample was applied on glow discharged carbon coated and formvar stabilized 400 mesh copper grids (Ted Pella) and incubated for approximately 30s. Excess sample was blotted off and the grid was washed with MilliQ water prior to negative staining using 2% uranyl acetate. TEM images were acquired using a Hitachi HT7700 (Hitachi High-technologies) transmission electron microscope operated at 100 kV equipped with a 2kx2k Veleta CCD camera (Olympus Soft Imaging System).

### Animal experiments

All animal experiments were performed in accordance with the ethical permits B4-16, B5-16 and Dnr. 20275-2021, and designed to minimize the suffering and pain of the animals. 10-weeks old, female C57BL/6 mice were used for the animal work.

For Nanoluc-mRNA *in vivo* experiments, equal amounts of Nanoluc-PUFe mRNA-loaded or control EVs were systemically injected into the mice intraperitoneally. The mRNA doses per injection were determined by the amount of actively loaded Nanoluc-PUFe mRNA per EV as measured by RT-qPCR. The corresponding EV doses ranged from 2.5×10^11^ EVs to 1×10^12^ EVs per animal. Mice were sacrificed at indicated time points and selected internal organs (liver, kidney, lung, spleen) were harvested. Organs were directly submerged in 1 ml lysis buffer (PBS supplemented with 0.01% TritonX-100), weighed, and lysed as described above to be analyzed using the NanoGlo luciferase assay.

For intratumoral delivery of EVs containing murine Ox40L-PUFe mRNA and protein, the B16F10 melanoma model was used. The tumors were established as previously described^61^. Briefly, female C57BL/6 (20 ± 2 g) mice were subcutaneously implanted with 7×10^5^ B16F10 cells on day 0. They were monitored daily for 6-10 days until they developed palpable tumors at sizes of 50-100 mm^3^. After tumor development, mice were treated by repetitive (every third day) intratumoral injections of particle-count matched mRNA-loaded or control EVs, or buffer only, in a total volume of 50 µl. The mice were observed daily, and tumor volumes were measured every third day. The mice were sacrificed if the tumor size exceeded 1,500 mm^3^ or if the mice met the scoring of pre-set humane end points (weight loss of more than 20% of initial weight and development of necrotic tumor tissue).

### Software

All illustrations were created using the BioRender online tool (publication licenses available upon request). Bar graphs and statistical analysis were done using GraphPad Prism software. Flow cytometry data was analyzed and visualized using the FlowJo Analysis Software (v 10.5.3 (FlowJo, LLC, BD Biosciences).

## ACKNOWLEDGEMENTS

We thank the Electron Microscopy Core Facility EMil at the Department of Laboratory Medicine, Karolinska Institutet, for the assistance in transmission electron microscopy.

## Additional Funding Sources

- European Research Council (ERC) under the European Union’s Horizon 2020 research and innovation programme (DELIVER, grant agreement No 101001374) (S.E.A.)
- European Union’s Horizon 2020 research and innovation programme (EXPERT, grant agreement No 825828) (S.E.A.)
- Swedish foundation of Strategic Research FormulaEx, SM19-0007 (S.E.A.)
- Cancerfonden project grant 21 1762 Pj 01 H (S.E.A.)
- Swedish Research Council grant 4–258/2021 (S.E.A.)
- Swedish Research Council grant 2021-02407 (J.N.)
- CIMED junior investigator grant (J.N.)

## AUTHOR CONTRIBUTIONS

S.E.A., J.N., J.H., D.G., and L.E. initiated this project. J.N., D.G., J.H., L.E. and X.L. designed the project and cloned the constructs. A.Z. and X.L. designed, performed, and analyzed most of the experiments. V.C.L., D.G., D.M., M.D.L., G.C., L.E., J.H., T.S., O.E., N.K., Z.N., G.Z., H.Z., S.R., O.W., A.G. contributed to design, performance, and analysis of experiments. S.E.A. and V.C.L were in charge of the overall project lead and management. A.Z. wrote the manuscript that was reviewed and complemented by all authors.

## COMPETING INTERESTS / INTELLECTUAL PROPERTY STATEMENT

The authors declare the following competing interests:

- S.E.A. is a co-founder, consultant, and stakeholder of Evox Therapeutics Ltd.
- J.N., D.G., and A.G. are consultants and stakeholders of Evox Therapeutics Ltd.
- V.C.L., M.D.L., L.E., J.H., and T.S. are current or former employees of Evox Therapeutics Ltd.
- This work is protected by patent families WO2019092145 and WO2020225392 owned by Evox Therapeutics Ltd. Patent information WO2019092145: applicant – Evox Therapeutics Ltd; inventors – J.N., D.G., L.E., and J.H.; international application number – PCT/EP2018/080681. Patent information WO2020225392: applicant – Evox Therapeutics Ltd; inventors – V.C.L, A.Z., X.L., M.D.L., and L.E.; international application number – PCT/EP2020/062791.
- O.E., G.C., N.K., Z.N., G.Z., H.Z., and S.R. declare no conflict of interest.

## DATA AVAILABILITY STATEMENT

All data are available in the main text, figures, or supplementary information. Primer sequences are listed in Table 1.

Construct sequences are available upon request.

## SUPPLEMENTARY FIGURES

**Figure S 1.**
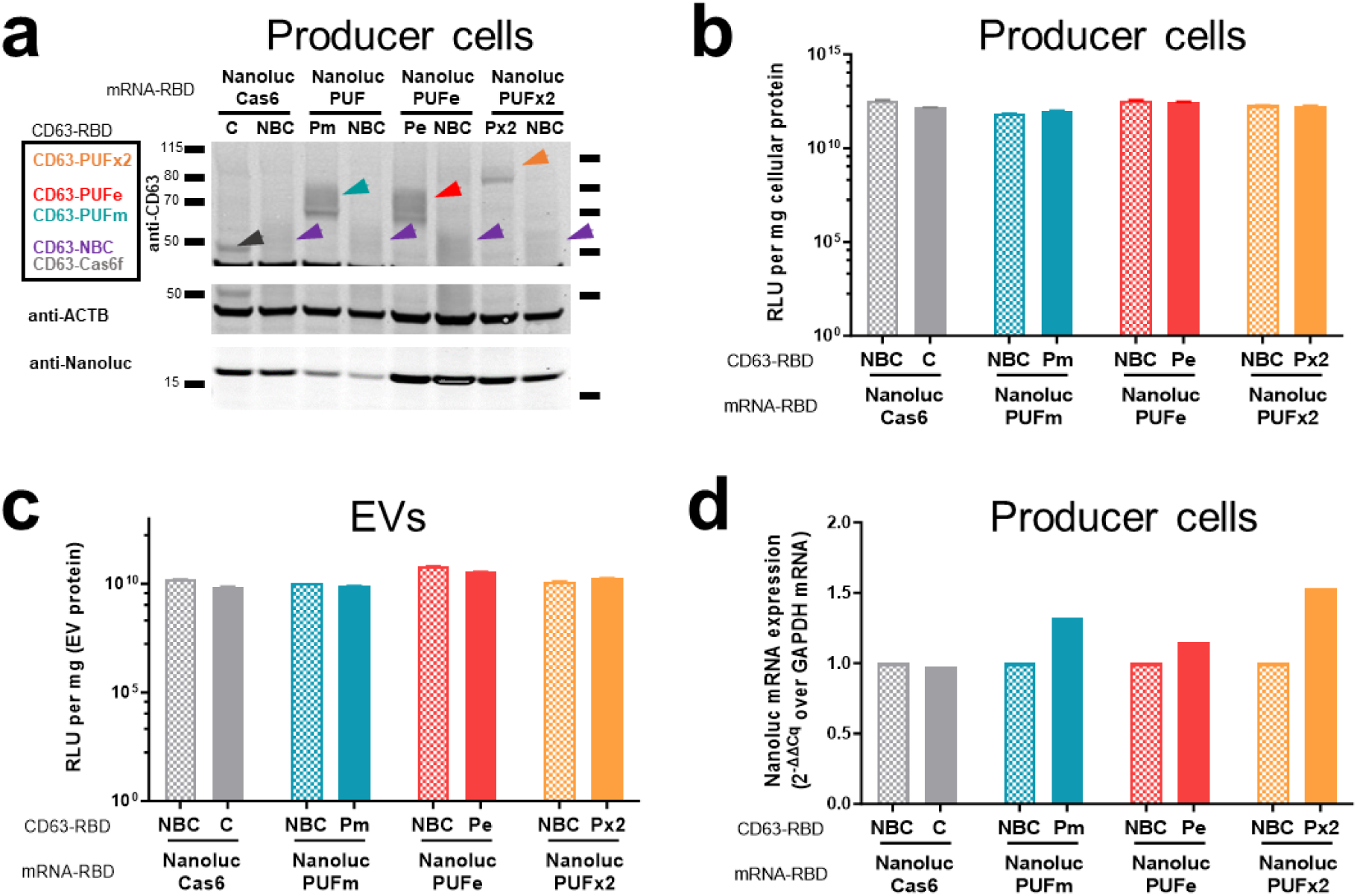
Characterization of mRNA stable EV producer cells and EVs. (a) Western Blot analysis of EVs produced from Nanoluc-RBD mRNA stable EV producer cells as indicated on top. The expression of the CD63-RBD fusion proteins, either CD63-Cas6f (C), CD63-PUFm (Pm), CD63-PUFe (Pe), CD63-PUFx2 (Px2), or CD63-NBC (NBC), respectively, was validated by probing for CD63, sizes are indicated with colored arrowheads. Moreover, Nanoluc protein expression was validated, ACTB expression served as reference. Protein loaded per lane: 30 µl cell lysate (4-15 µg total protein) (b) and (c) Nanoglo luminescence assay to determine the amount of active Nanoluc enzyme per mg (c) total cellular protein and (d) EV protein. The respective Nanoluc-RBD mRNA is indicated below, CD63-RBD fusion protein expression above: CD63-Cas6f (C), CD63-PUFm (Pm), CD63-PUFe (Pe), CD63-PUFx2 (Px2), or CD63-NBC (NBC). (d) Relative quantification by RT-qPCR of Nanoluc mRNA normalized on GAPDH reference mRNA in Nanoluc-RBD (RBD motif as indicated) mRNA stable EV producer cells transfected for EV production.

**Figure S 2.**
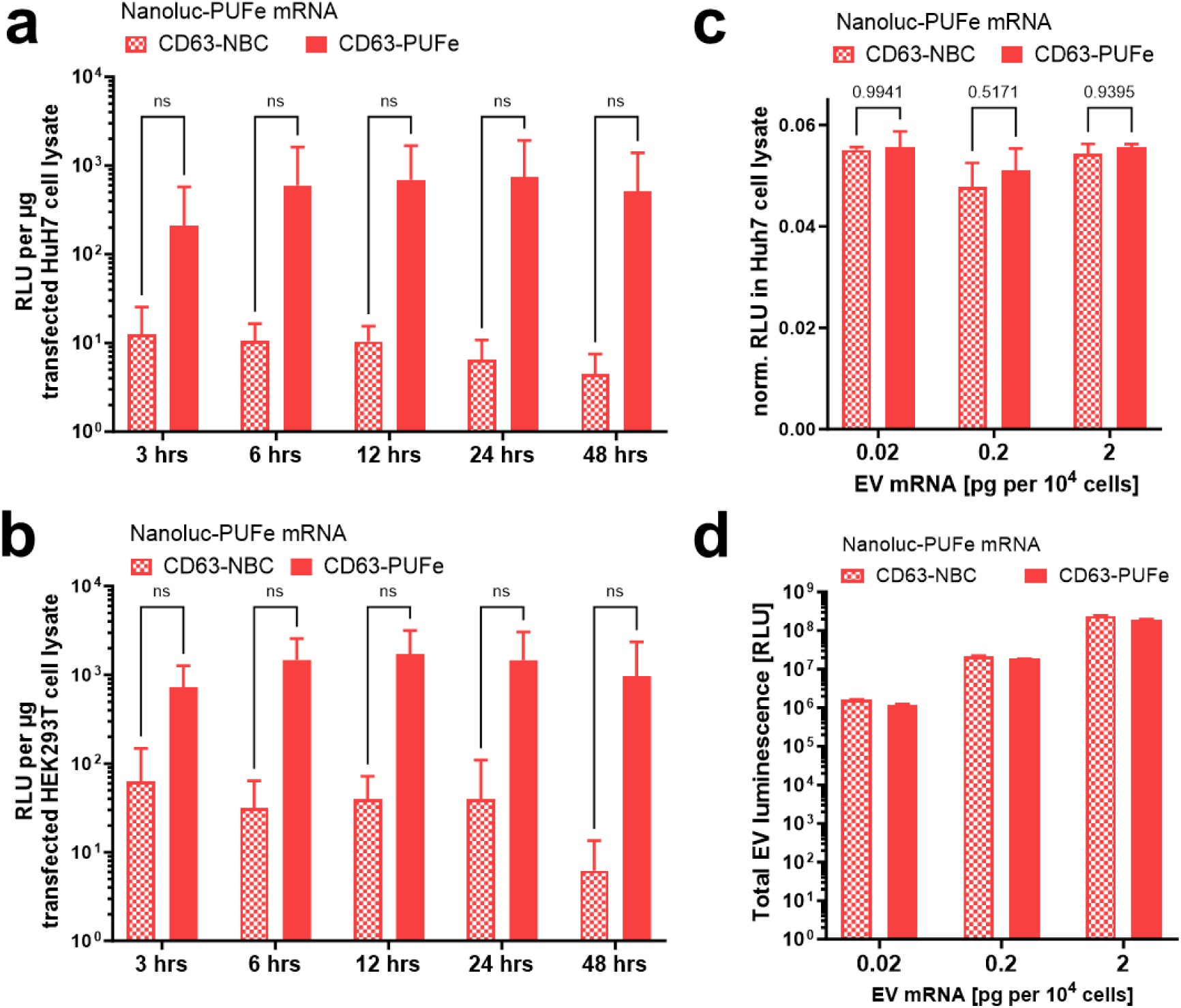
EV mRNA cargo is functional and delivered *in vitro*. (a) and (b) Purified EV RNA from Nanoluc-PUFe mRNA EVs and control EVs was transfected into (a) Huh7 cells or (b) HEK293T cells and Nanoluc protein activity was measured at indicated timepoints. RNA from mRNA EVs gave rise to high Nanoluc protein expression, proving functionality of EV-derived engineered mRNA in recipient cells. Experiment were performed with n=3 and statistically analyzed by Two-Way ANOVA. (c) Uptake of Nanoluc-PUFe mRNA EVs at increasing mRNA doses (0.02 pg, 0.2 pg, and 2 pg Nanoluc mRNA per 1×10^4^ cells) or particle count-matched control EVs in Huh7 recipient cells. Cellular Nanoluc protein activity was measured at 24 h and normalized to Nanoluc protein measured in EVs. Increased ratio for cells treated with Nanoluc-PUFe mRNA EVs compared to cells treated with control EVs indicated translation. Experiment were performed with n=3 and statistically analyzed by Two-Way ANOVA. (d) Exemplified Nanoluc protein content as measured in relative luminescence (RLU) of different amounts of EVs used for treatment *in vitro*. These values were used to normalize Nanoluc protein expression in lysates of treated cells, as they accounted for the co-delivered passively loaded protein without replenishment by mRNA translation.

**Figure S 3.**
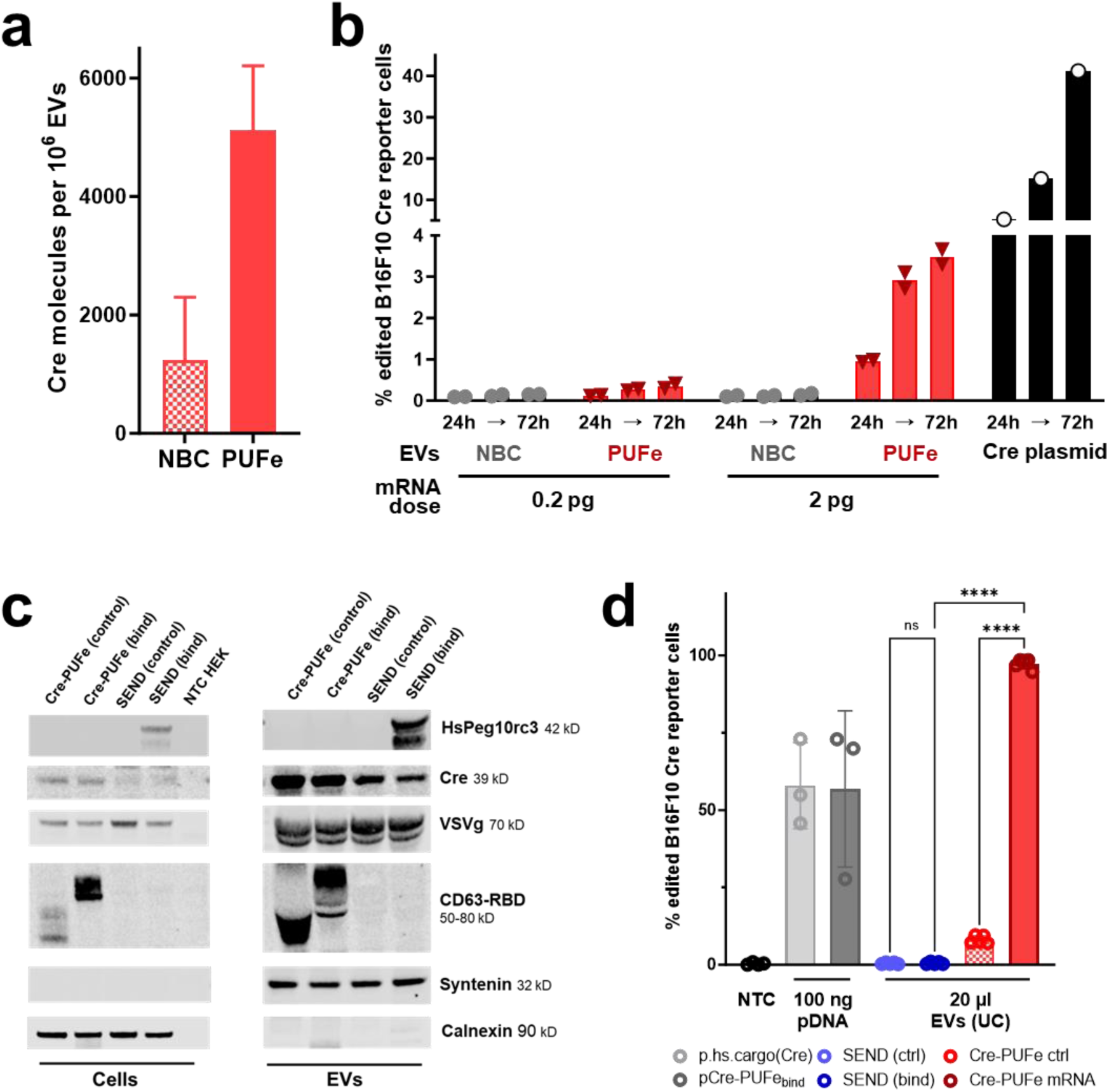
EV-mediated delivery of Cre recombinase *in vitro*. (a) Absolute quantification of Cre-PUFe mRNA loading per 1×10^6^ EVs in control (NBC) and mRNA EVs (PUFe). (b) In vitro uptake analysis of B16F10 Traffic Light Cre reporter cells treated with equal amounts of Cre-PUFe mRNA EVs at a dose of 0.2 pg and 2 pg mRNA per 1×10^4^ cells (red), or particle count-matched control EVs (grey), both with VSVg expression. Frequencies of genomically edited Cre reporter cells were assessed at 24, 48, and 72 hours post treatment by flow cytometry. Cells treated with mRNA loaded EVs show a time- and dose-dependent increase in genomically edited cell frequencies. Cre plasmid transfection (black) served as positive control. Significance level α=0.05, Two-Way ANOVA. (c) Western Blot analysis to validate HsPEG10rc3, Cre, VSVg, and CD63-PUFe or CD63-NBC expression in EV producer cells (left) and EVs (right). Syntenin and Calnexin expression served as reference for EVs and cells, respectively. Cell lysates: 14 µg protein per lane, EVs: 123 µg per lane. (d) *In vitro* uptake analysis of B16F10 Traffic Light Cre reporter cells treated with 20 µl of EV suspension prepared by ultracentrifugation. Cre-PUFe (control) or (bind) – CD63-PUFe platform (red), SEND (control) or (bind) - selective endogenous encapsidation for cellular delivery platform (Segel et al., Science, 2021) with or without HsPeg10rc3 expression (blue), 100 ng plasmid transfection encoding Cre mRNA with PUFe or hsPeg10rc3 binding sequences (grey) served as positive control.

**Figure S 4.**
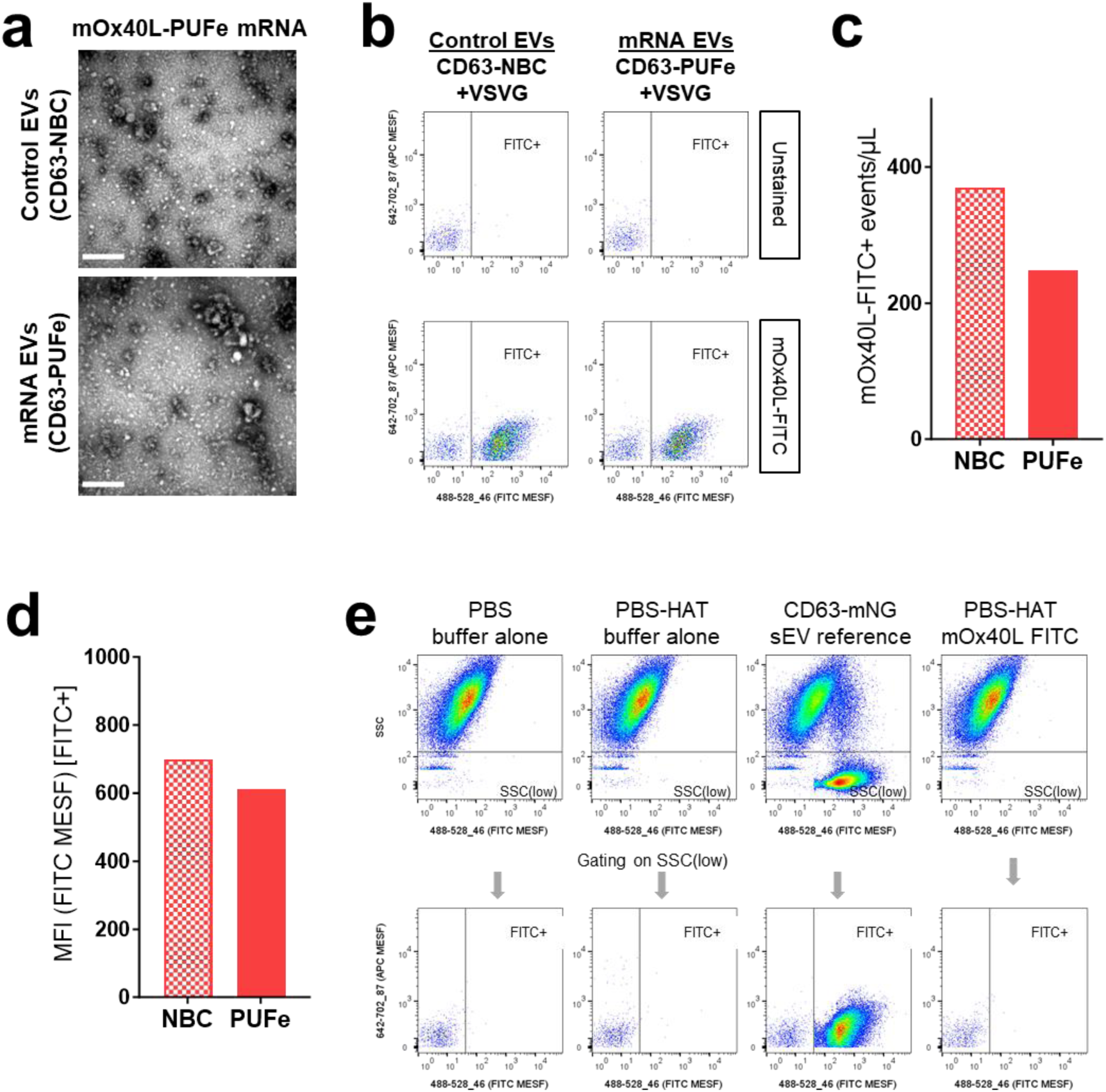
Characterization of mOx40L-PUFe mRNA EVs. (a) Negative stain Transmission Electron Microscopy images of control EVs and mRNA EVs loaded with mOx40L-PUFe mRNA. Scale bar: 300 nm (b) SSC(low)-gated data from control EVs and mRNA EVs loaded with mOx40L-PUFe mRNA, either unstained or stained with anti-mouse OX40L-FITC antibodies. Dotplots show FITC signals (excitation laser: 488 nm; emission filter: 528/46 nm) in FITC MESF units versus autofluorescence (APC channel; excitation laser: 642 nm; emission filter: 702/87 nm) in APC MESF units. (c) Quantification of the concentration of mOx40L-FITC positive events as measured post staining and dilution. (d) Quantification of the mean fluorescence intensity (MFI) of FITC+ gated events in FITC MESF units. (e) Buffer controls, antibody control, and pre-gating strategy for identification of SSC(low) events equivalent to small EVs (sEVs) based on previously established workflows and CD63-mNG green fluorescent reference sEVs.

**Figure S 5.**
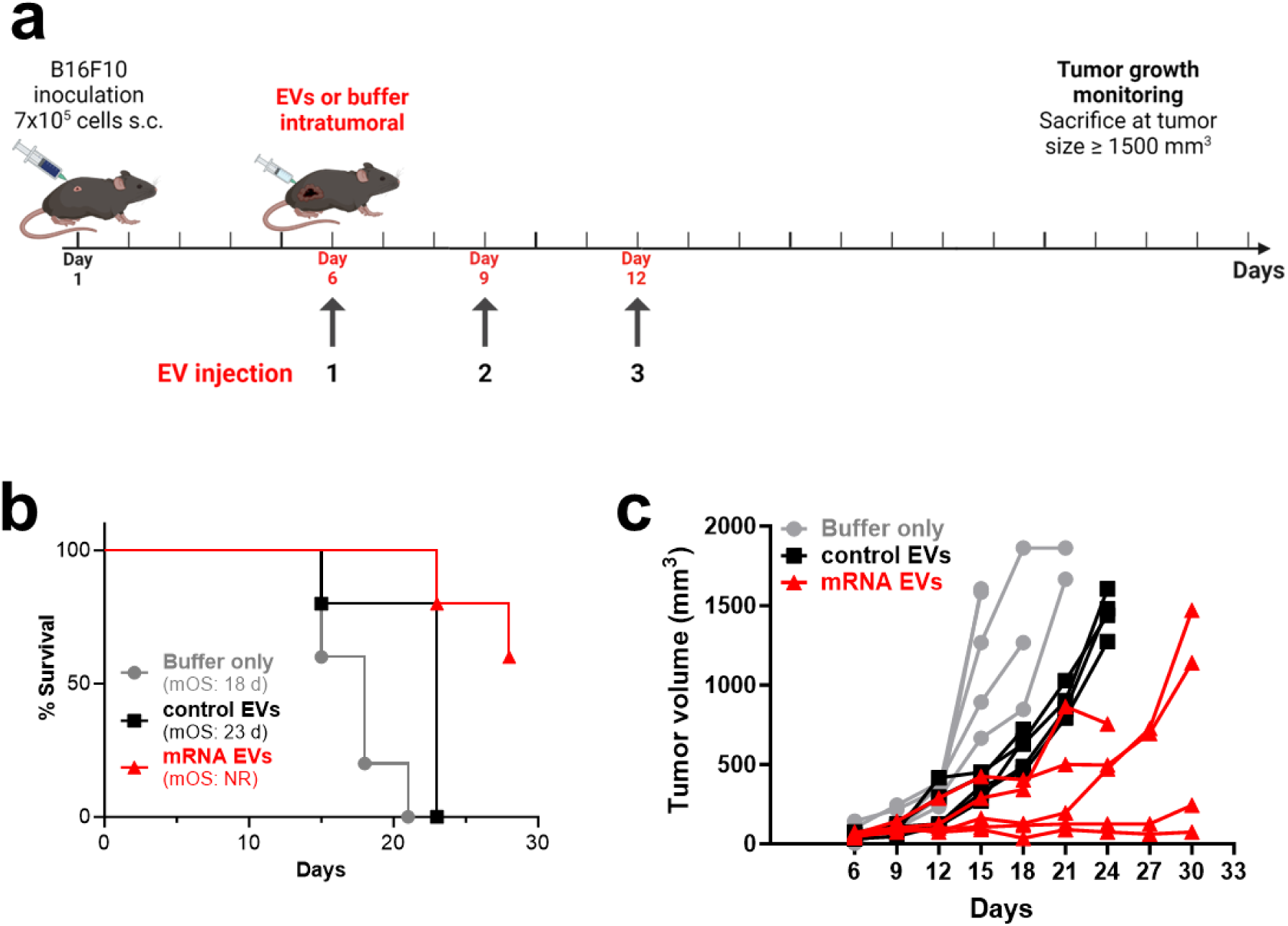
Efficient EV-mediated delivery of the immunomodulatory molecule mOx40L in murine tumor model *in vivo*. **(a)** Injection scheme for intratumoral injections of mRNA EVs, control EVs, or suspension buffer (PBS-HAT) at a dose of 2 ng mRNA per kg bodyweight into B16F10 melanoma-bearing mice (n=5 per group). After tumor engraftment, mice were injected 3 times and monitored regularly for tumor growth. Figure created using BioRender. (b) Kaplan-Meier survival analysis of mice treated with mOx40L mRNA EVs, control EVs, or buffer only with assessment of median overall survival (mOS). All curves are significantly different from each other (Log-rank (Mantel-Cox) test, P value 0.0006. NR – not registered. (c) Tumor volumes measured regularly after the last injection. Each line represents one mouse of the respective group. Two out of five of the mRNA EV-treated mice went into complete remission and lost their tumor beyond palpability for the duration of the experiment (30 days). Curve analysis (Wilcoxon Signed Rank Test): Buffer only group P value (two-tailed) 0.0312, Control EV group P value (two-tailed) 0.0156, mRNA EV group P value (two-tailed) 0.0039, α=0.05, all curves significant.

## Supporting information

Full blot images

