## Supplementary material for "Novel Endogenous Engineering Platform for Robust Loading and Delivery of Functional mRNA by Extracellular Vesicles"

### SUPPLEMENTARY FIGURES

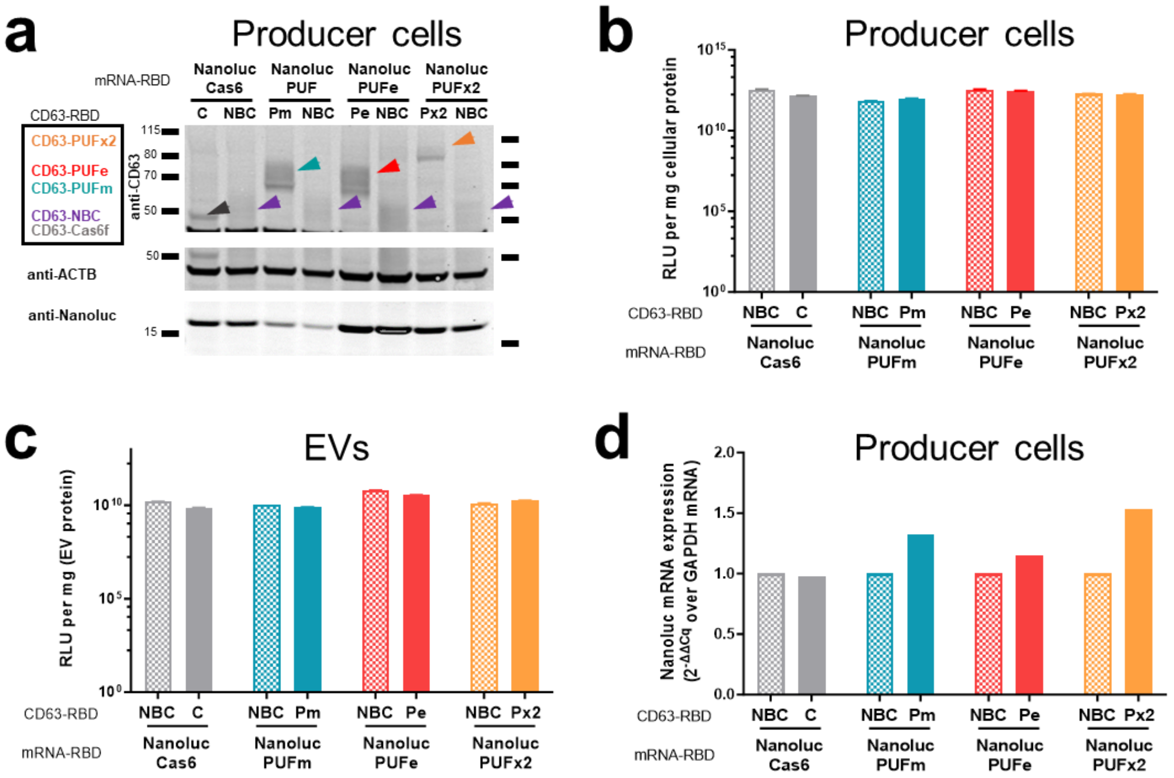


**Figure S 1. Characterization of mRNA stable EV producer cells and EVs. (a)** Western Blot analysis of EVs produced from Nanoluc-RBD mRNA stable EV producer cells as indicated on top. The expression of the CD63-RBD fusion proteins, either CD63-Cas6f (C), CD63-PUFm (Pm), CD63-PUFe (Pe), CD63-PUFx2 (Px2), or CD63-NBC (NBC), respectively, was validated by probing for CD63, sizes are indicated with colored arrowheads. Moreover, Nanoluc protein expression was validated, ACTB expression served as reference. Protein loaded per lane: 30 µl cell lysate (4-15 µg total protein) **(b) and (c)** Nanoglo luminescence assay to determine the amount of active Nanoluc enzyme per mg (c) total cellular protein and (d) EV protein. The respective Nanoluc-RBD mRNA is indicated below, CD63-RBD fusion protein expression above: CD63-Cas6f (C), CD63-PUFm (Pm), CD63-PUFe (Pe), CD63-PUFx2 (Px2), or CD63-NBC (NBC). **(d)** Relative quantification by RT-qPCR of Nanoluc mRNA normalized on GAPDH reference mRNA in Nanoluc-RBD (RBD motif as indicated) mRNA stable EV producer cells transfected for EV production.

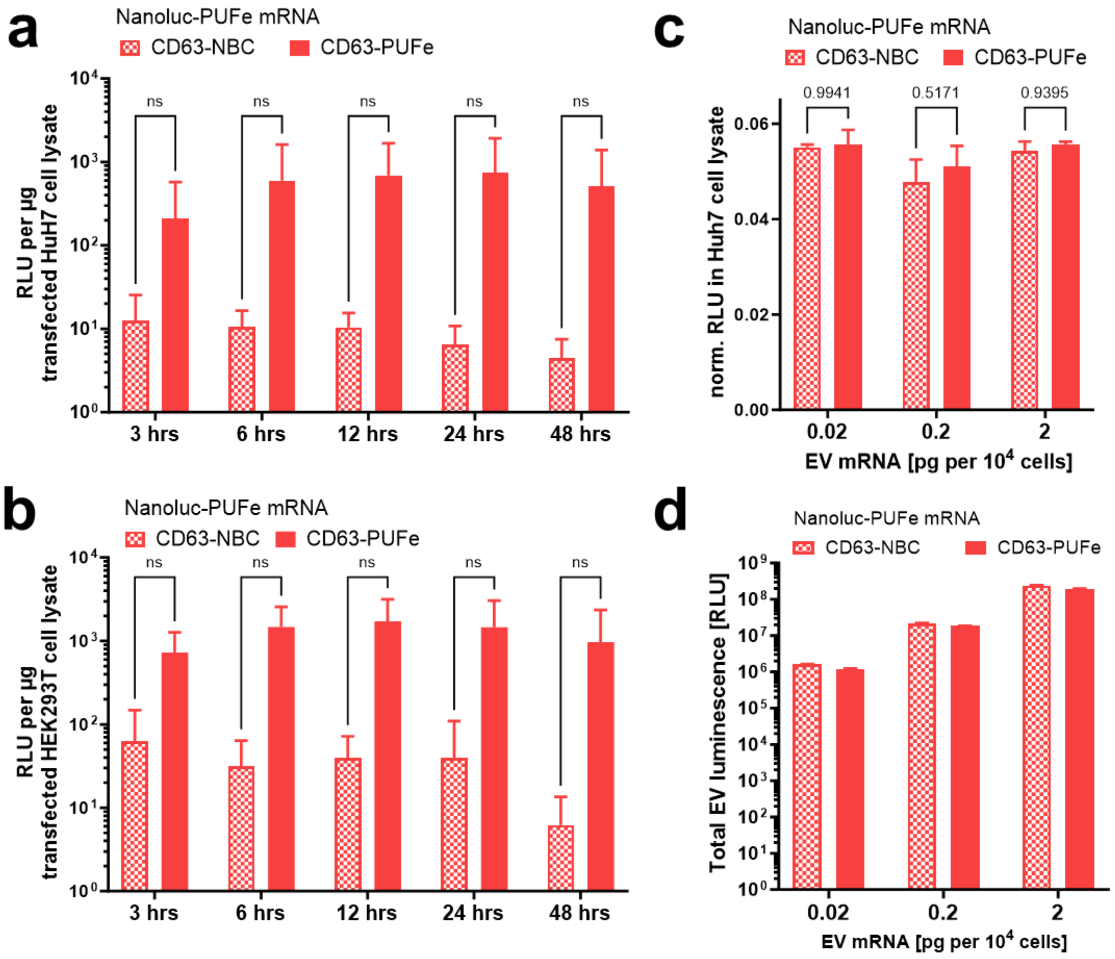


**Figure S 2. EV mRNA cargo is functional and delivered *in vitro*.** (a) and (b) Purified EV RNA from Nanoluc-PUFe mRNA EVs and control EVs was transfected into (a) Huh7 cells or (b) HEK293T cells and Nanoluc protein activity was measured at indicated timepoints. RNA from mRNA EVs gave rise to high Nanoluc protein expression, proving functionality of EV-derived engineered mRNA in recipient cells. Experiment were performed with n=3 and statistically analyzed by Two-Way ANOVA. (c) Uptake of Nanoluc-PUFe mRNA EVs at increasing mRNA doses (0.02 pg, 0.2 pg, and 2 pg Nanoluc mRNA per $1x10^4$ cells) or particle count-matched control EVs in Huh7 recipient cells. Cellular Nanoluc protein activity was measured at 24 h and normalized to Nanoluc protein measured in EVs. Increased ratio for cells treated with Nanoluc-PUFe mRNA EVs compared to cells treated with control EVs indicated translation. Experiment were performed with n=3 and statistically analyzed by Two-Way ANOVA. (d) Exemplified Nanoluc protein content as measured in relative luminescence (RLU) of different amounts of EVs used for treatment *in vitro*. These values were used to normalize Nanoluc protein expression in lysates of treated cells, as they accounted for the co-delivered passively loaded protein without replenishment by mRNA translation.

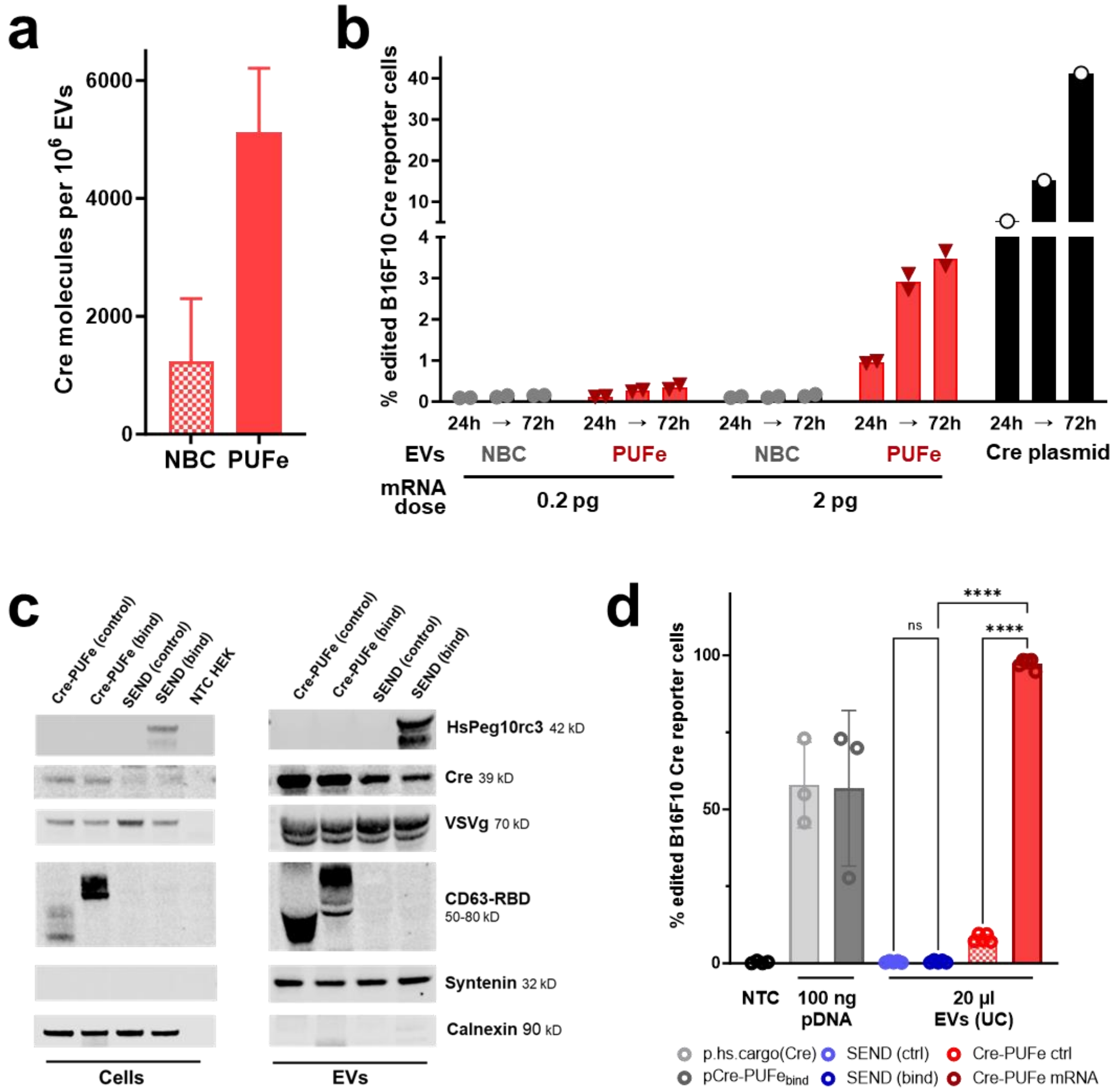


**Figure S 3. EV-mediated delivery of Cre recombinase *in vitro*.** (a) Absolute quantification of Cre-PUFe mRNA loading per $1x10^6$ EVs in control (NBC) and mRNA EVs (PUFe). (b) *In vitro* uptake analysis of B16F10 Traffic Light Cre reporter cells treated with equal amounts of Cre-PUFe mRNA EVs at a dose of 0.2 pg and 2 pg mRNA per $1x10^4$ cells (red), or particle count-matched control EVs (grey), both with VSVg expression. Frequencies of genomically edited Cre reporter cells were assessed at 24, 48, and 72 hours post treatment by flow cytometry. Cells treated with mRNA loaded EVs show a time- and dose-dependent increase in genomically edited cell frequencies. Cre plasmid transfection (black) served as positive control. Significance level α=0.05, Two-Way ANOVA. (c) Western Blot analysis to validate HsPEG10rc3, Cre, VSVg, and CD63-PUFe or CD63-NBC expression in EV producer cells (left) and EVs (right). Syntenin and Calnexin expression served as reference for EVs and cells, respectively. Cell lysates: 14 µg protein per lane, EVs: 123 µg per lane. (d) *In vitro* uptake analysis of B16F10 Traffic Light Cre reporter cells treated with 20 µl of EV suspension prepared by ultracentrifugation. Cre-PUFe (control) or (bind) – CD63-PUFe platform (red), SEND (control) or (bind) - selective endogenous encapsidation for cellular delivery platform (Segel et al., Science, 2021) with or without HsPeg10rc3 expression (blue), 100 ng plasmid transfection encoding Cre mRNA with PUFe or hsPeg10rc3 binding sequences (grey) served as positive control.

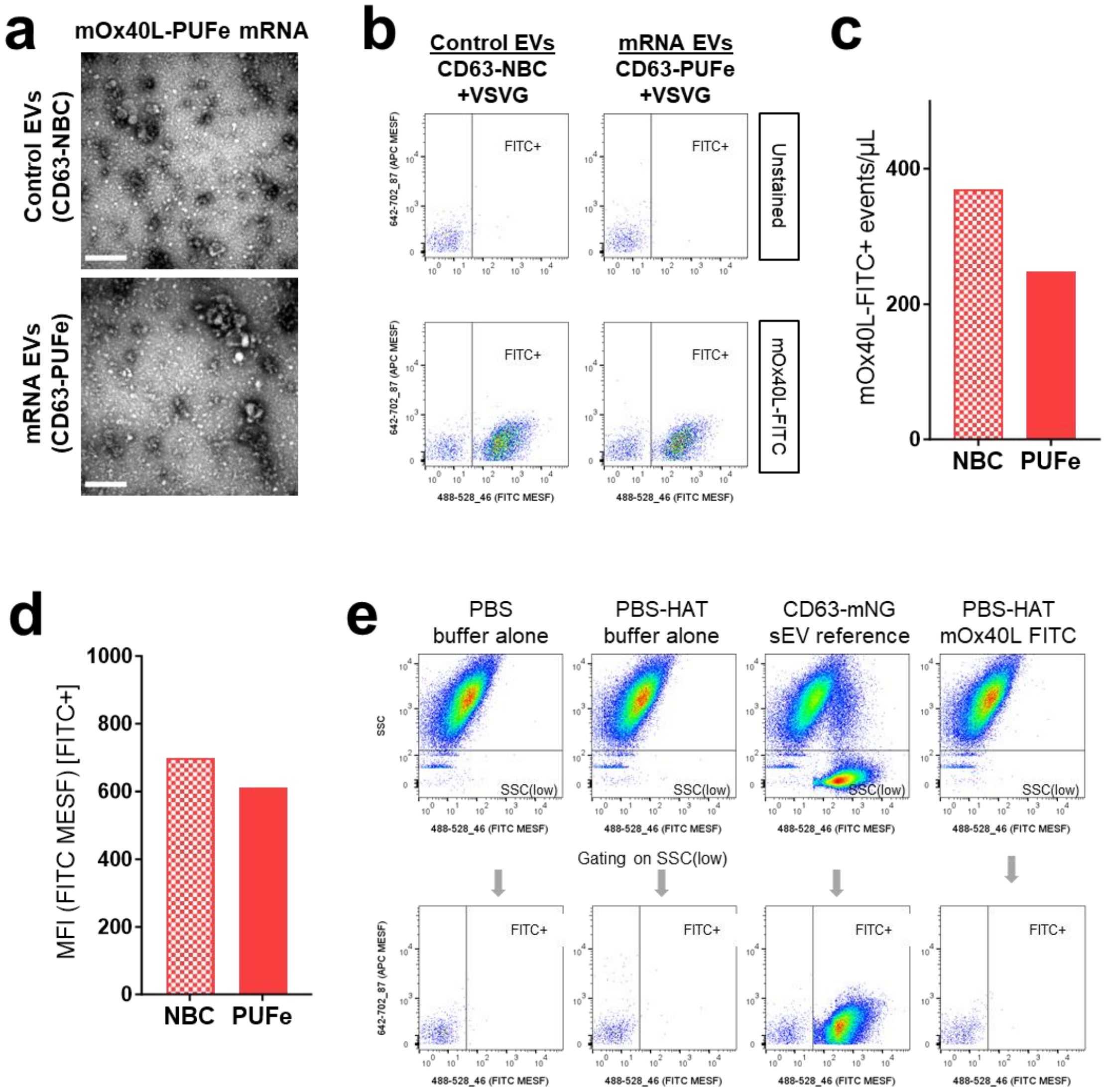


**Figure S 4. Characterization of mOx40L-PUFe mRNA EVs.** (a) Negative stain Transmission Electron Microscopy images of control EVs and mRNA EVs loaded with mOx40L-PUFe mRNA. Scale bar: 300 nm (b) SSC(low)-gated data from control EVs and mRNA EVs loaded with mOx40L-PUFe mRNA, either unstained or stained with anti-mouse OX40L-FITC antibodies. Dotplots show FITC signals (excitation laser: 488 nm; emission filter: 528/46 nm) in FITC MESF units versus autofluorescence (APC channel; excitation laser: 642 nm; emission filter: 702/87 nm) in APC MESF units. (c) Quantification of the concentration of mOx40L-FITC positive events as measured post staining and dilution. (d) Quantification of the mean fluorescence intensity (MFI) of FITC+ gated events in FITC MESF units. (e) Buffer controls, antibody control, and pre-gating strategy for identification of SSC(low) events equivalent to small EVs (sEVs) based on previously established workflows and CD63-mNG green fluorescent reference sEVs.

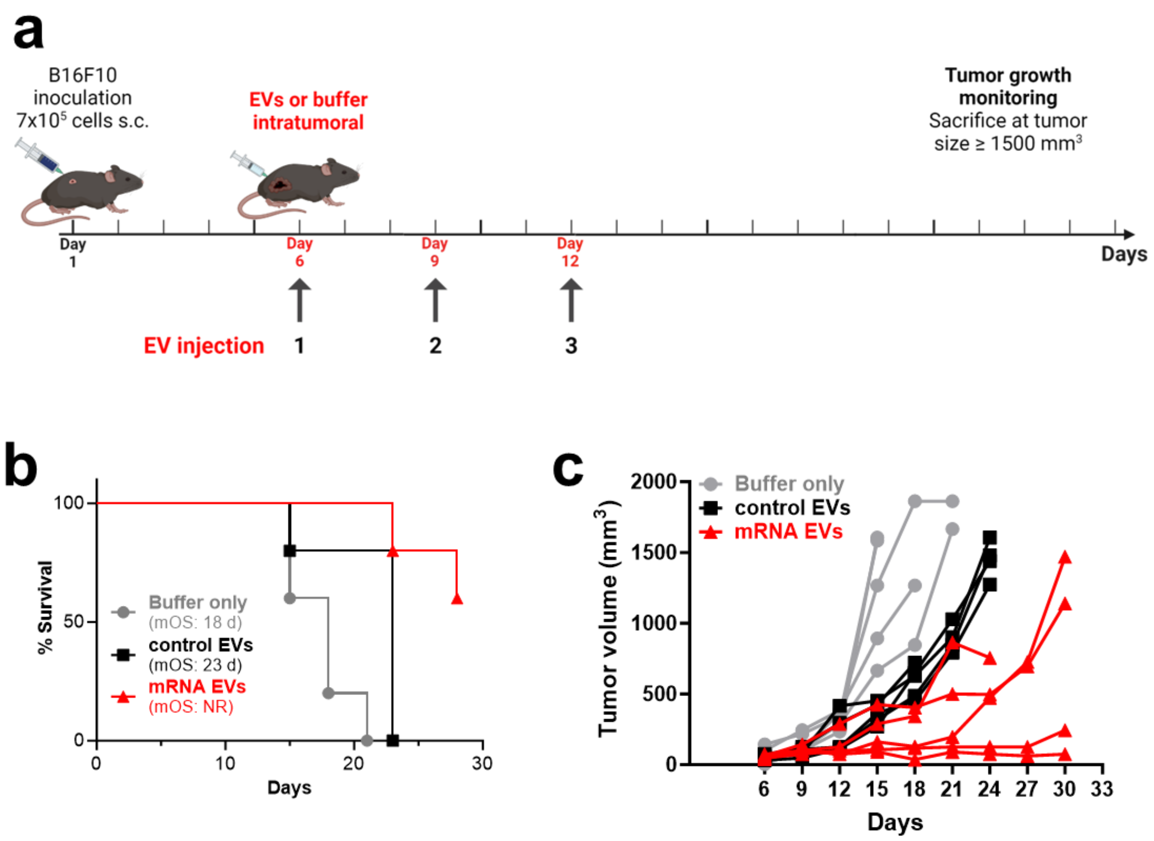


**Figure S 5. Efficient EV-mediated delivery of the immunomodulatory molecule mOx40L in murine tumor model *in vivo*. (a**) Injection scheme for intratumoral injections of mRNA EVs, control EVs, or suspension buffer (PBS-HAT) at a dose of 2 ng mRNA per kg bodyweight into B16F10 melanoma-bearing mice (n=5 per group). After tumor engraftment, mice were injected 3 times and monitored regularly for tumor growth. Figure created using BioRender. (b) Kaplan-Meier survival analysis of mice treated with mOx40L mRNA EVs, control EVs, or buffer only with assessment of median overall survival (mOS). All curves are significantly different from each other (Log-rank (Mantel-Cox) test, P value 0.0006. NR – not registered. (c) Tumor volumes measured regularly after the last injection. Each line represents one mouse of the respective group. Two out of five of the mRNA EV-treated mice went into complete remission and lost their tumor beyond palpability for the duration of the experiment (30 days). Curve analysis (Wilcoxon Signed Rank Test): Buffer only group P value (two-tailed) 0.0312, Control EV group P value (two-tailed) 0.0156, mRNA EV group P value (two-tailed) 0.0039, α=0.05, all curves significant.

**FULL BLOT IMAGES**

Full Blot Image Figure 1d

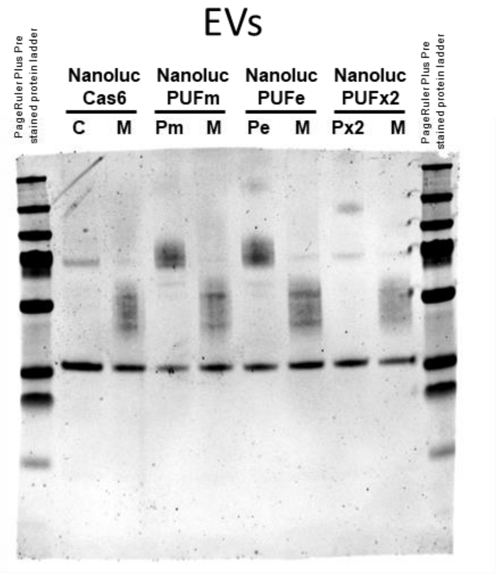


Primary antibodies:

- rb-α-CD63 (ab68418, Abcam, 1:1000)
- ms-α-SDCBP (Syntenin) (TA504796, Thermo Fisher Scientific, 1:500)

Full Blot Images Figure 3c

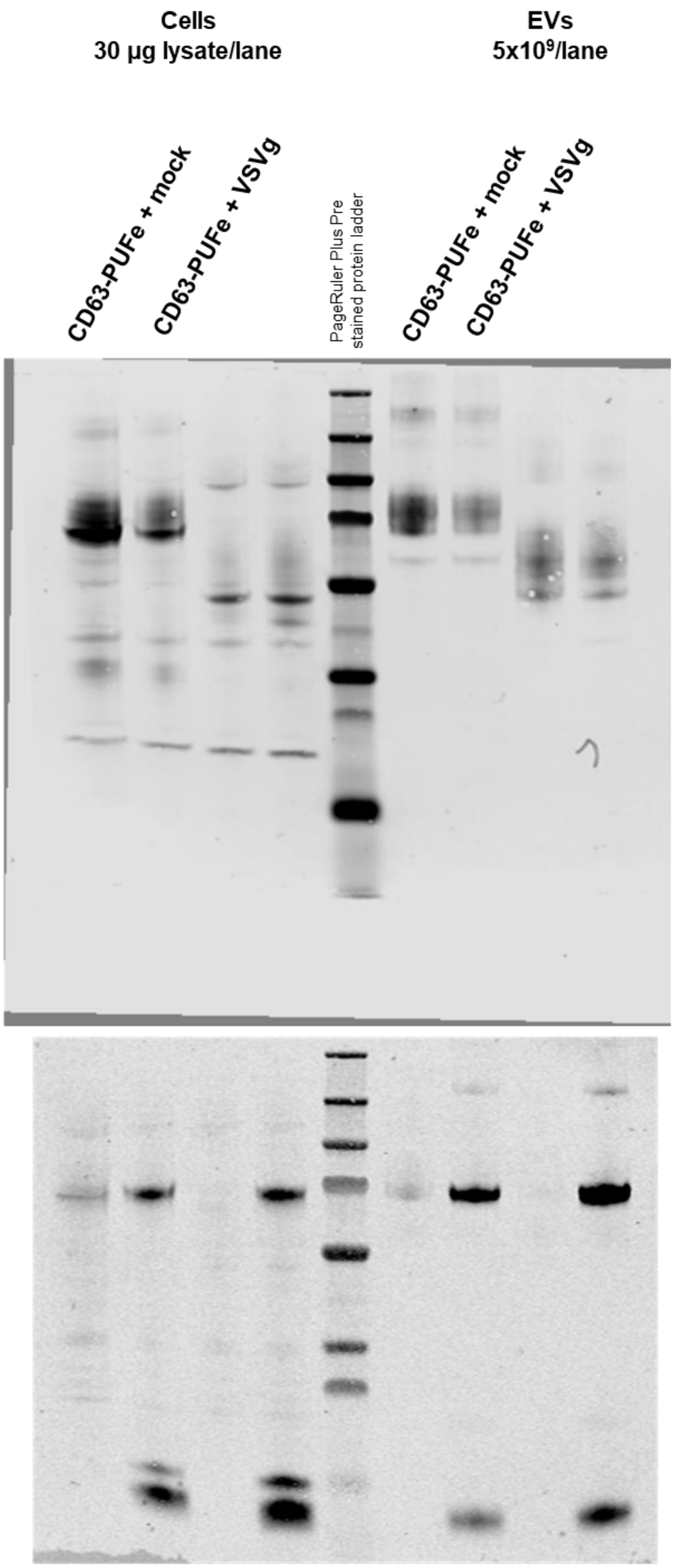


Primary antibody:
- rb-α-CD63 (ab68418, Abcam, 1:1000)

Primary antibody:
- gt-α-VSV-G Tag (PA1-30278, Thermo Fisher Scientific, 1:1000)

Full Blot Images Supplementary Figure S 1a

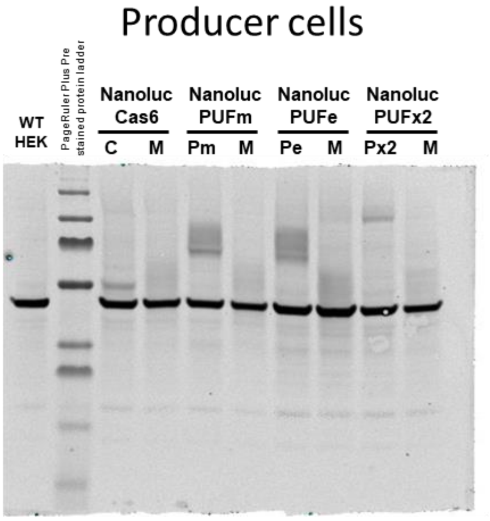


Primary antibodies:
- rb-α-CD63 (ab68418, Abcam, 1:1000)
- ms-α-ACTB (A5441, Merck, 1:20000)

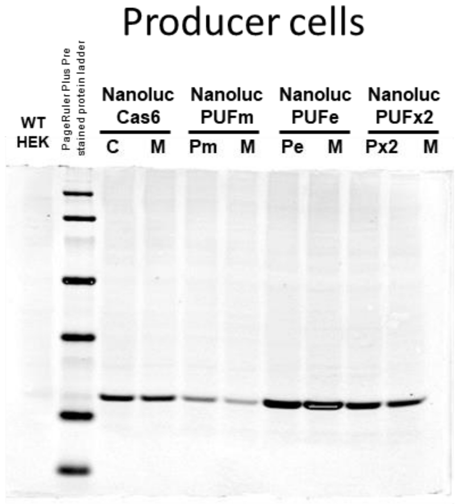


Primary antibody:
- rb-α-Nanoluc (non-commercial, Promega, 1:1000)

Full Blot Images Supplementary Figure S 3c

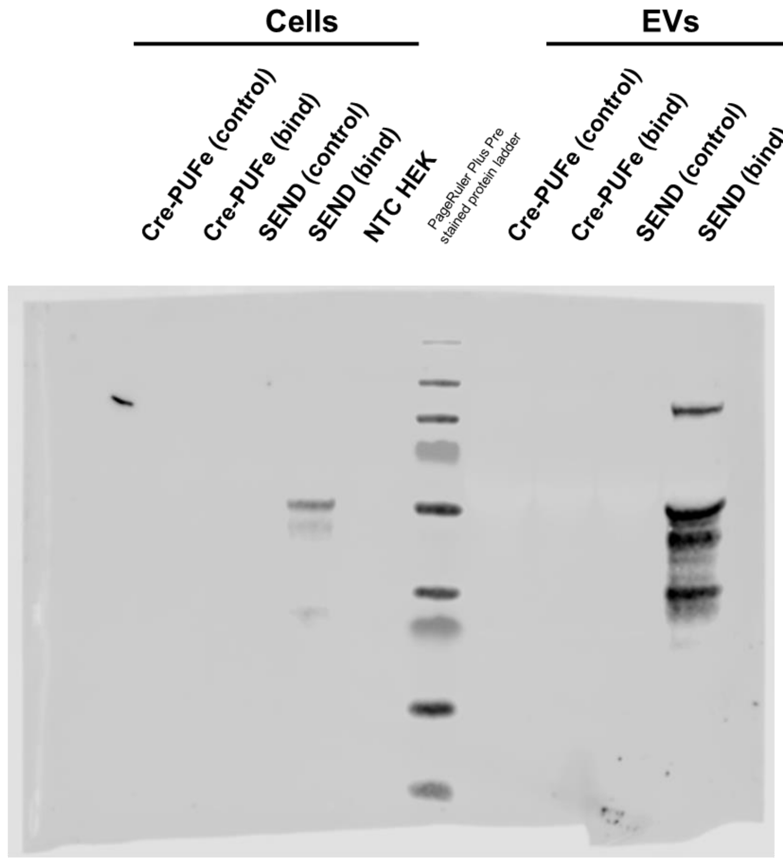


Primary antibody:
- rb-α-PEG10 (#77111, Cell Signaling Technology, 1:1000)

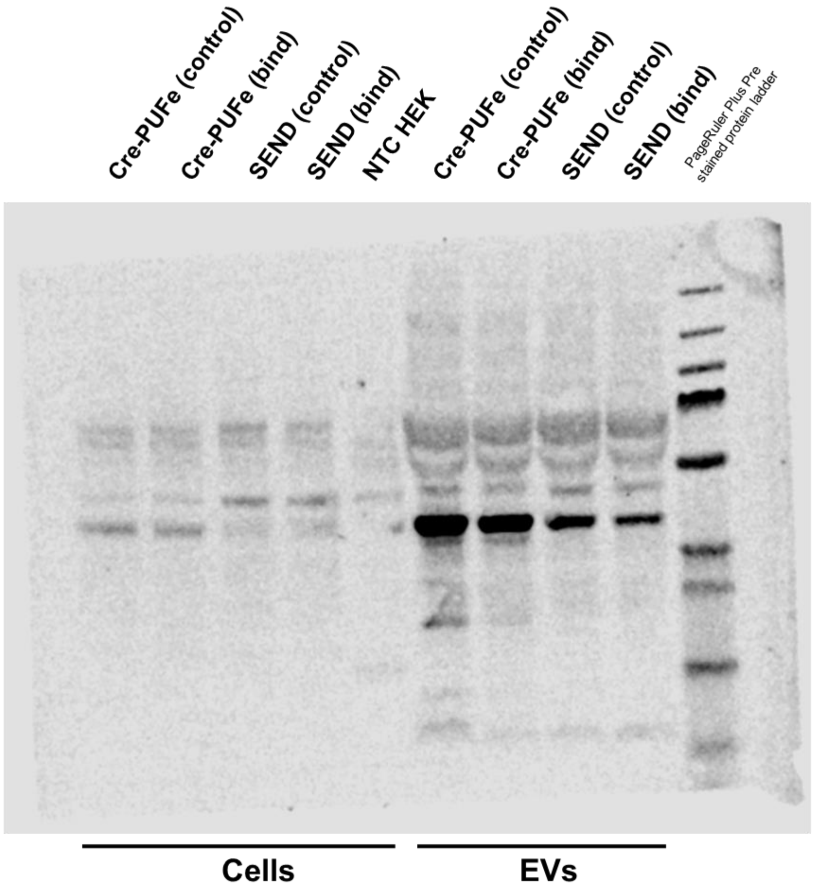


Primary antibody:
- rb-α-Cre recombinase (ab188568, Abcam, 1:1000)

Full Blot Images Supplementary Figure S 3c

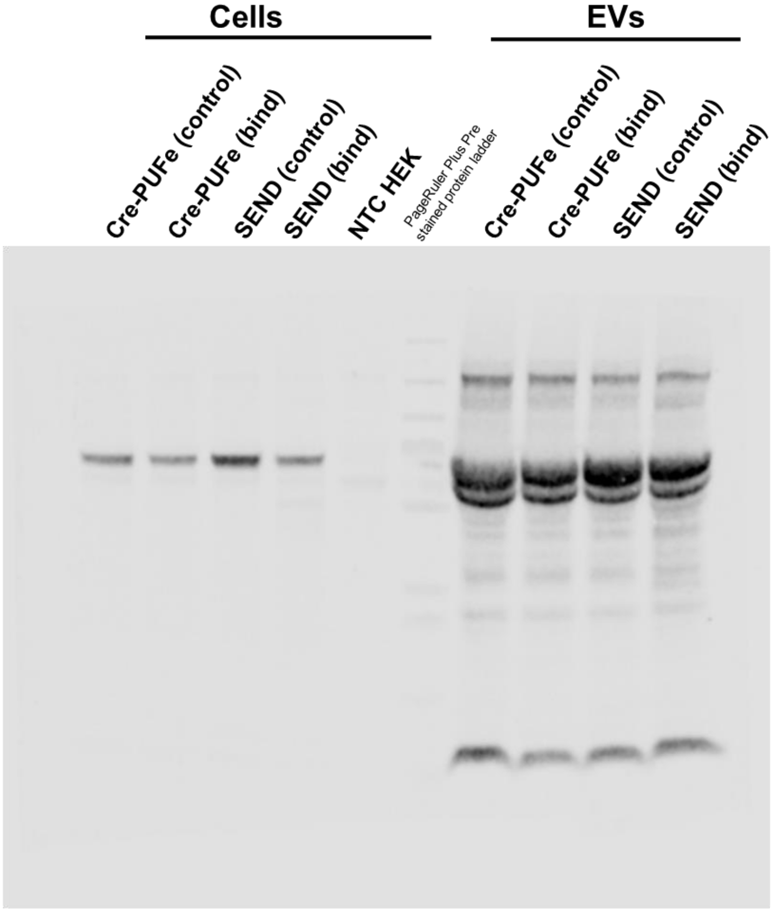


Primary antibody:

- gt-α-VSV-G Tag (PA1-30278, Thermo Fisher Scientific, 1:1000)

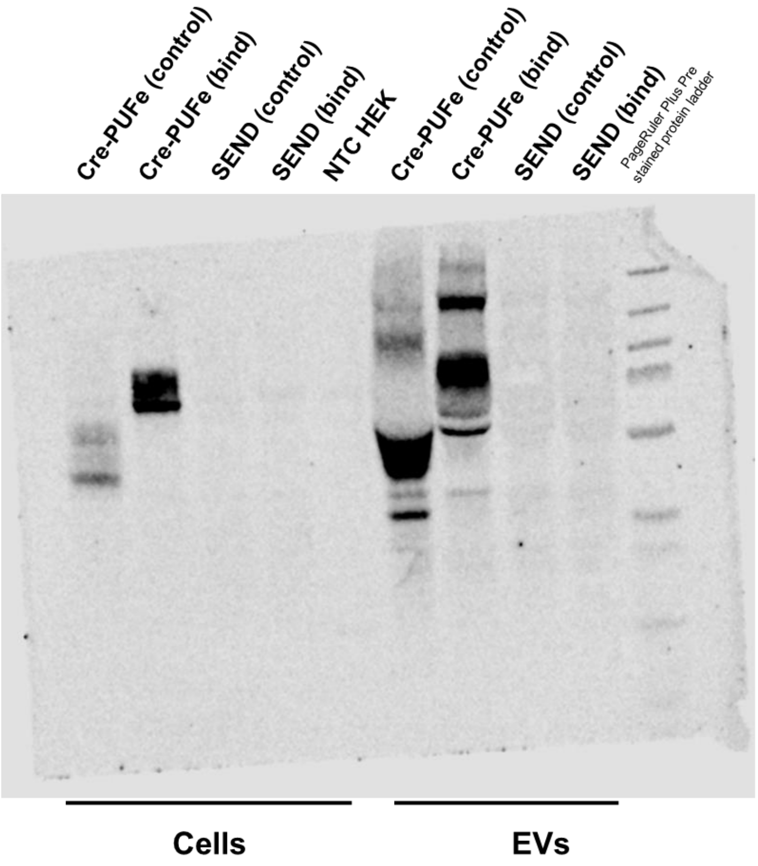


Primary antibody:

- rb-α-CD63 (ab68418, Abcam, 1:1000)

Full Blot Images Supplementary Figure S 3c

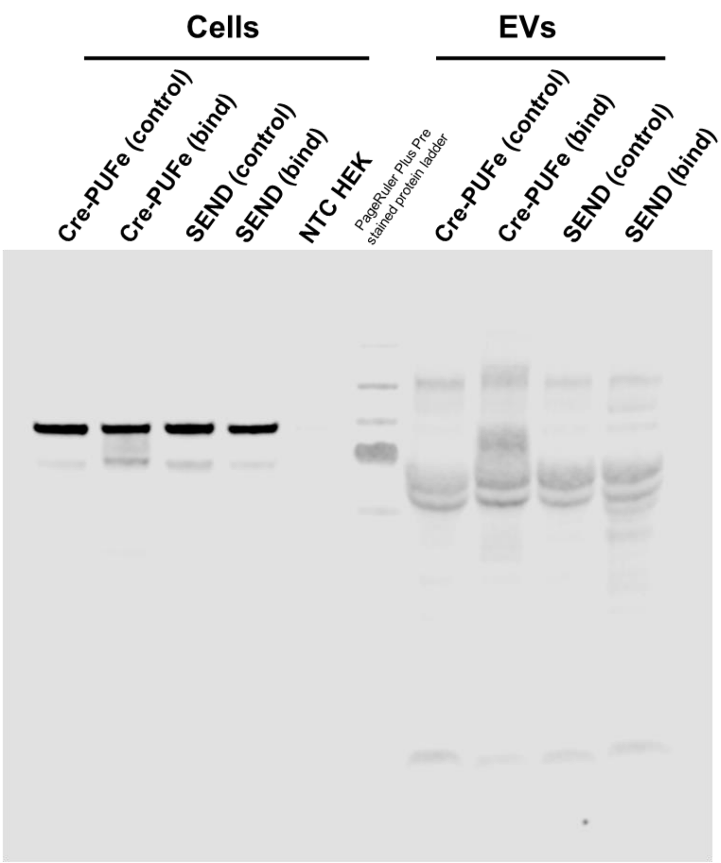


Primary antibody:

- rb-α-Calnexin (ab22595, Abcam, 1:1000)

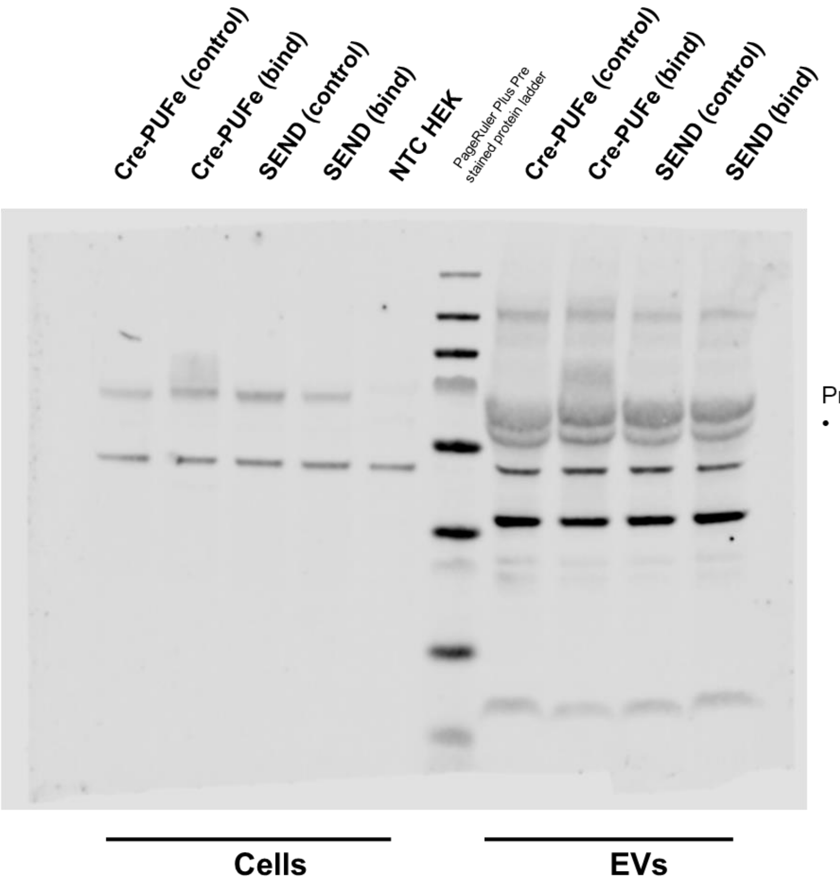


Primary antibody:

- ms-α-SDCBP (Syntenin) (TA504796, Thermo Fisher Scientific, 1:500)
